# Tissue mechanics regulate mitotic nuclear dynamics during epithelial development

**DOI:** 10.1101/688374

**Authors:** Natalie J. Kirkland, Alice C. Yuen, Melda Tozluoglu, Nancy Hui, Ewa K. Paluch, Yanlan Mao

## Abstract

Cell divisions are essential for tissue growth. In pseudostratified epithelia, where nuclei are staggered across the tissue, each nucleus migrates apically before undergoing mitosis. Successful apical nuclear migration is critical to preserve tissue integrity during cell division. Most previous investigations have focused on the local cellular mechanisms controlling nuclear migration. Yet, inter-species and inter-organ comparisons of different pseudostratified epithelia suggest global tissue architecture may influence nuclear dynamics, but the underlying mechanisms remain elusive. Here, we use the developing *Drosophila* wing disc to systematically investigate, in a single epithelial type, how changes in tissue architecture during growth influence mitotic nuclear migration. We observe distinct nuclear dynamics at discrete developmental stages, as epithelial morphology changes. We then use genetic and physical perturbations to show a direct effect of cell density on mitotic nuclear positioning. We also find Rho kinase and Diaphanous, which facilitate mitotic cell rounding in confined cell conditions, are essential for efficient apical nuclear movement. Strikingly, perturbation of Diaphanous causes increasing defects in apical nuclear migration as the tissue grows, and these defects can be reversed by acute physical reduction of cell density. Our findings reveal how the mechanical environment imposed on cells within a tissue alters the molecular and cellular mechanisms adopted by single cells for mitosis. We speculate that mechanical regulation of apical mitotic positioning could be a global mechanism for tissue growth control.

## Introduction

Successful mitosis in densely packed epithelia relies on the ability of cells to round up. Rounding is driven by reorganisation of cellular actin into a highly contractile actomyosin cortex at the cell membrane [1–6] and generates the physical space required for mitotic spindle assembly and chromosome separation [2, 7]. Indeed, mitotic rounding is required to resist the mechanical constraints exerted by neighbouring cells and prevent aberrations or failure of mitosis [3, 8]. In most epithelia, mitotic cells round up at the apical surface of the tissue, facilitating planar-orientated cell divisions [9, 10]. Maintaining cell divisions in the plane of the epithelium preserves mono-layered epithelial organisation and tissue integrity, and prevents cancerous delamination of cells [9, 11].

Pseudostratified epithelia (PSE), where nuclei are staggered across an epithelial monolayer, present amongst the most densely packed tissue organisation. PSE are found in abundant and diverse developing tissues, from the intestine [12] to the neural tube [13] and the neocortex [14]. Accordingly, the architecture of PSE can differ drastically, varying in height, nuclear composition and curvature [15]. For instance, PSE can range from 20 *µ*m to several millimetres (reviewed by [16]). Similarly, the number of nuclei staggered across the monolayer can vary, as can nuclear density [15]. The often extensive height and nuclear layering, warrant that for division to occur apically, many PSE must translocate nuclei to the apical surface prior to mitosis, a process referred to as interkinetic nuclear migration (IKNM) [17].

Investigations of IKNM have revealed a variety of mechanisms driving nuclear movement (reviewed in [15, 16, 18]). Tall neuroepithelia found in the developing cortex utilise microtubule-dependent mechanisms [19–23], while shorter neuroepithelia, including the zebrafish retina and hindbrain utilise actomyosin-dependent mechanisms [24–27]. In both cases, apical nuclear movement occurs in the G2 stage of the cell cycle, prior to mitotic rounding [20-22, 24, 25]. However, examples of *Drosophila* tissue of similar height to the zebrafish retina and hindbrain, primarily the wing imaginal disc [26] and the short, pseudostratified neuroepithelia [27], appear to drive nuclear movement coincident with mitotic cell rounding, a mechanism conserved from sea anemone, *Nemostella vectensis* [26].

Even though previous studies have mostly focused on the processes controlling IKNM locally, in individual cells, comparative studies of PSE suggest that IKNM dynamics might also be influenced by tissue architecture. Comparison of IKNM in the mouse and ferret cortex, show a reduced rate of apical motion in denser and stiffer tissue [28, 29]. Furthermore, faster apical nuclear movements are observed in the developing zebrafish hindbrain compared to the retina, which differ in tissue shape [25]. The retina exhibits positive curvature at the apical surface, whereas the hindbrain is flat. Mechanical constraints exerted by the epithelial morphology may give rise to the observed differences in nuclear dynamics. This idea is supported by evidence for mechanical regulation of cell differentiation in the developing zebrafish neural tube, where gradual apical nuclear crowding alters the positioning of progenitor nuclei and thus their exposure to signalling molecules [30]. To what extent, and how tissue properties influence nuclear dynamics remains unclear, as currently, only epithelia from different species have been directly compared [28, 29].

Here, we use the *Drosophila* wing disc to investigate the effect of tissue architecture on IKNM in a single epithelium as it grows and changes in morphology. The larval wing disc is a monolayered epithelial structure comprised largely of a pseudostratified, columnar layer that increases in size until transforming into the adult fly wing. We find that mitotic nuclear dynamics change over the time course of wing disc development, coincident with changes in tissue height, nuclear layer composition and cell density. We then use genetic and mechanical perturbations to alter cell density and find a marked effect on apical mitotic positioning, indicative of altered mitotic nuclear dynamics. Finally, we investigate the role of Rho kinase (Rok) and the formin Diaphanous (Dia), two key drivers of mitotic rounding [2–4, 8], in IKNM. We show that while Rok is indispensible for efficient apical nuclear movement and mitotic positioning at all developmental stages, the dependency on Dia increases as development, and thus cell density, increases. Our findings reveal how the mechanical environment imposed on cells confined within a tissue can influence molecular and cellular mechanisms.

## Results

### Wing disc development is associated with increased tissue height, nuclear layering and cell density

To identify features of cell and tissue morphology that may influence mitotic nuclear behaviour, we first characterized how the apico-basal architecture of the larval wing disc epithelium changes during development. We focused on wing discs at 72, 96 and 120 hours after egg laying (AEL) as they exhibit distinct tissue morphologies. Wing discs at 72 hours AEL are mostly flat, while discs at 96 and 120 hours AEL display three characteristic tissue folds but have clear differences in their projected surface areas between the two time points [31, 32] (Figure 1A). We fixed wing discs at the three selected developmental stages and stained for actin, nuclei and mitotic cells, using phalloidin, DAPI and anti-phospho histone H3 (PH3) staining, respectively. We then quantified epithelial architecture in the pouch region of the wing disc (Figures 1A, S1A). We found at 72 hours AEL, the apico-basal height of the pouch epithelium measured 20.75 ± 1.38 *µ*m, increasing to approximately 28.02 ± 1.95 *µ*m at 96 hours AEL and finally 38.79 ± 3.09 *µ*m by 120 hours AEL (Figures 1A,A’,B’ and S1B). We observed that the increasing epithelia height also accompanied differences in the nuclear layer organisation along the apico-basal axis (Figure 1A’). To quantitatively assess nuclear organisation along the apico-basal axis, we measured the heights of three characteristic zones along this axis: the apical proliferative zone, the nuclear layer and the basal nucleus-free zone (Figures 1A’-B’ and S1A-E). The apical proliferative zone was defined as a region largely devoid of interphase nuclei, but occupied by mitotic cells rounded at the apical surface. The nuclear layer comprised the region directly below the apical proliferative zone, where interphase nuclei adopt staggered positions, characteristic of PSE. Finally, the basal nucleus-free zone was the region below the nuclear layer devoid of all nuclei. We found that the most striking change in height through development was displayed by the nuclear layer, which started at a height of 10.95 ± 1.01 *µ*m at 72 hours AEL, increasing to 16.91 ± 1.409 *µ*m at 96 hours AEL and finally 22.26 ± 1.97 *µ*m at 120 hours AEL (Figures 1A’,B’ and S1D). We also found that the average number of nuclei stacked across the nuclear layer, as well as nuclei density, increased as development progressed (Figures 1C,D). This is consistent with the suggestion for the vertebrate cortex, that nuclear layering increases when cell number increases faster than the accommodating surface area [33].

**Figure 1.**
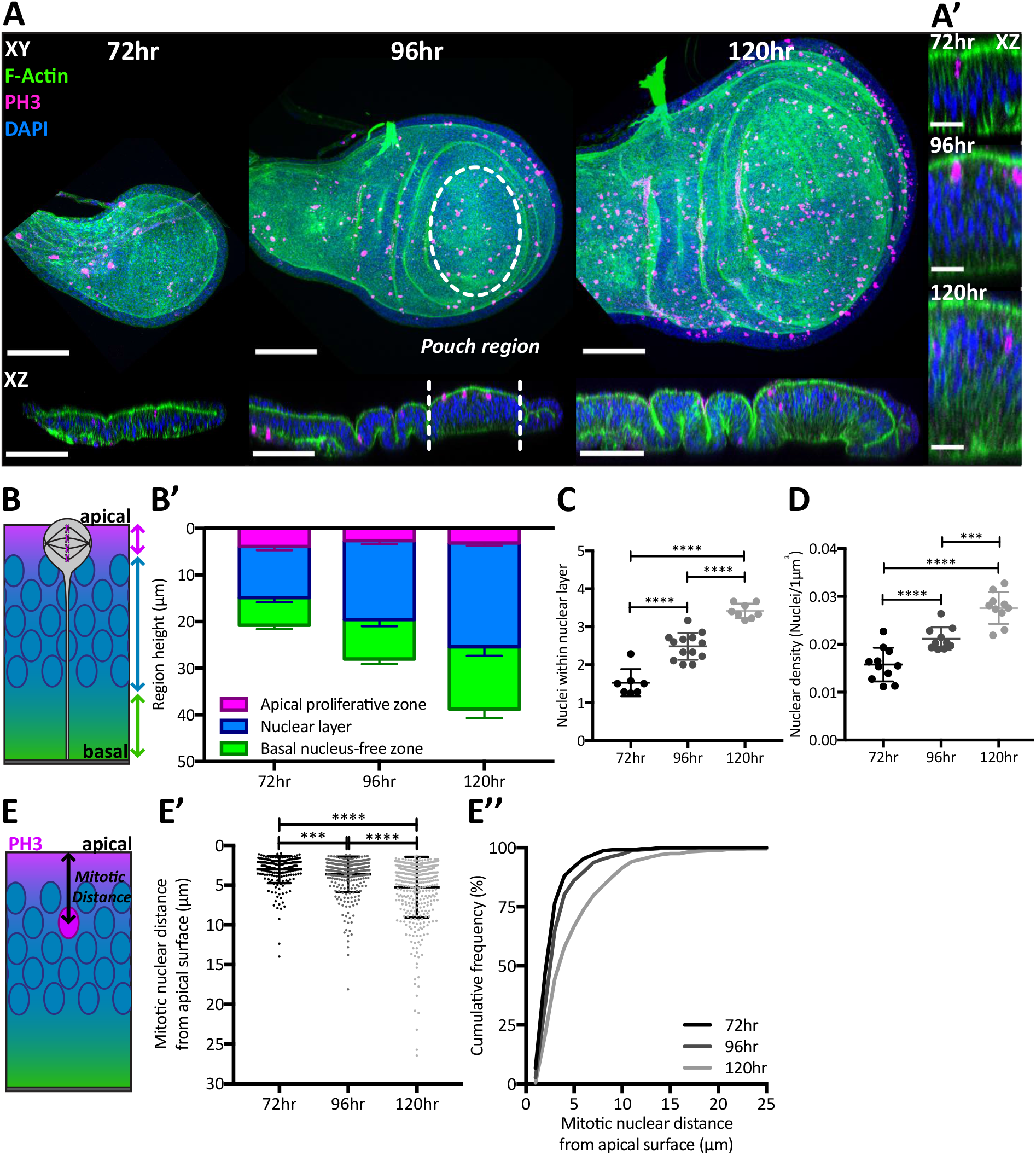
Changes in the wing disc architecture during development are associated with distinct patterns of mitotic nuclear positioning. (A) Drosophila wing imaginal disc at 72, 96 and 120 hours after egg laying (AEL) stained with phalloidin (green), anti-PH3 (magenta) and DAPI (blue). Top: projection images, bottom: single lateral cross sections through the centre of the wing disc (anterior-posterior axis). Wing disc pouch region highlighted with white dashed lines. (A’) Enlarged lateral cross section of the pouch region. (B) Schematic of lateral cross section of the pouch region, highlighting sub-regions along the apico-basal axis quantified in B’. Magenta represents the apical proliferative zone, blue represents the nuclear layer and green represents the basal nucleus-free zone. (B’) Quantification of height for the apico-basal sub-regions through development in the pouch region of the wing disc. Measurements acquired as described in Figure S1A. Break down of each segment presented in Figure S1B-E. (C) Number of nuclei within the nuclear layer region through development in the pouch region. Each dot represents the average of a single disc and acquired as described in Figure S1A. (D) Density of nuclei per 1*µ*m3 in the pouch region through development. (E) Schematic showing measurement of mitotic nuclear distance by measuring the distance of PH3+ nuclei from the apical surface, marked with F-Actin. (E’) PH3+ nuclei distance, referred to as mitotic nuclear distance, from apical surface in the pouch region of the wing disc at developmental stages presented as a dot plot. (E’’) Data in E’ presented as a cumulative frequency distribution. Normalised data of E shown in Figure S1G. (B’ and E’-E’’) n = 8, 8 and 5 wing discs for 72, 96 and 120hr respectively. (C) n = 7, 13 and 8 wing discs for 72, 96 and 120hr respectively. (D) n = 3 wing discs per stage. Scale bars: A, 50*µ*m, A’, 10*µ*m. Statistical significance in C-D are given for one-way ANOVA. Statistical significance in E given for Kolmogorov-Smirnov comparison of cumulative distribution. ***p < 0.001, ****p < 0.0001. Error bars represent mean ± SD in all plots.

Together, our findings demonstrate that during the development of the wing disc, the layering of nuclei increases and nuclei pack closer together to accommodate the increasing cell number accompanying growth.

### Developmental changes in tissue architecture are associated with distinct patterns of mitotic nuclear positioning

We then asked whether mitotic nuclear behaviour differed in discs with different PSE architectures. Previous work in the wing disc has proposed that apical positioning of the mitotic nucleus occurs during mitosis, by observing mitotic nuclei at, or close to the apical surface, and their mislocalisation away from the apical surface upon perturbation of mitotic rounding [26]. We therefore used PH3 immunostaining to mark mitotic nuclei and measured the distance from the centre of PH3+ nuclei to the apical surface at the distinct developmental stages, in the pouch region of the discs (Figure 1E). While the majority of PH3+ nuclei were within a few microns of the apical surface, we found that as development progressed, more PH3+ nuclei were positioned further from the apical surface (Figures 1E’,E’’). We observed non-apical PH3+ nuclei exhibited centrosome arrangements reminiscent of early mitosis (Figure S1F). As metaphase always occurs at the apical surface in wild type wing disc, this indicates that our fixed tissue analysis captures cells at different stages of IKNM. The developmental differences in the length of the nuclear layers, and their position within the epithelium, impacts the potential distance a mitotic nucleus may travel during IKNM. To account for these differences we normalised each distance to the combined height of the apical proliferative zone and the nuclear layer (the “nuclear region”, see Figure S1G). Upon normalisation, we found that mitotic nuclei were the most apically distributed at 96 hours AEL compared to wing discs at 72 and 120 hours AEL (Figures S1G’ and S1G’’).

Together, our results indicate that differences in epithelial morphology that accompany wing disc development may influence the distribution of mitotic nuclei captured in fixed tissues.

### Directed apical nuclear movement accompanies mitosis

The broader distribution of mitotic nuclei from the apical surface as development progresses suggests there could be differences in mitotic nuclear dynamics (Figures 1E-E’’ and S1G-G’’). In order to examine whether there is a developmental influence on mitotic nuclear dynamics, we first verified that apical movement is in fact driven during mitosis in the wing disc PSE. We live-imaged wing discs expressing Non-muscle Myosin-II-GFP (Sqh-GFP) and Histone 2A-RFP (His-RFP) and manually tracked nuclei by measuring the distance from the basal side of the nucleus to the apical surface of the wing pouch at each developmental stage (Figure 2). At all developmental stages, visual assessment of individual and averaged nuclear trajectories suggested nuclei moved randomly until a sharp transition to apically directed movement occurred shortly before metaphase (Figures 2A,A’ and S2A-C). In our analysis of the nuclei trajectories, we found that the average instantaneous nuclear velocities initially fluctuated around 0, followed by a persistent increase in instantaneous nuclear velocity for all developmental stages (Figures 2B, 2C and 2D). The transition to apical, directed movement correlated with enrichment of Sqh-GFP at the cell cortex (Figures 2F-F’’, white arrows) and progression to apical mitotic rounding (Figures 2F-F’’).

**Figure 2.**
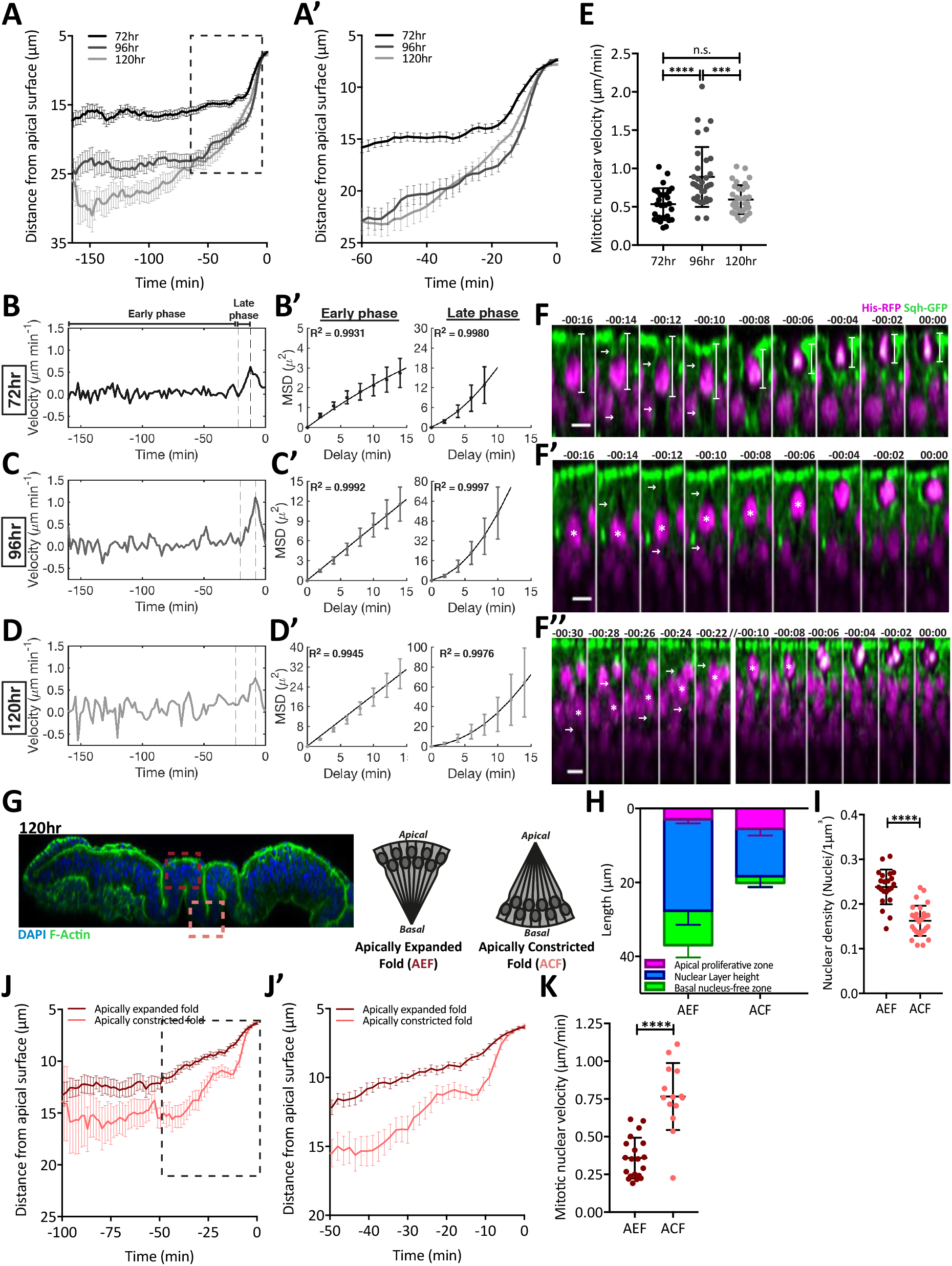
Mitotic nuclear dynamics depend on tissue architecture. (A-A’) Averaged tracks of mitotic nuclei within the wing disc pouch region at 72, 96 and 120 hours AEL, obtained by measuring distances illustrated in F, 72hr. 0 minutes is defined as the final metaphase nucleus. Black dotted line highlights inset shown in A’ enlarging region presenting the transition to directed apical movement. All tracks are presented in Figure S2A-C (B-D) Average instantaneous velocity measurements for tracked nuclei in A-A’. ‘Early phase’ denotes region showing non-directed movement as confirmed by mean square displacement analysis in B’-D’, left panel. ‘Late phase’ denotes region with directed nuclear movement as confirmed by mean square displacement analysis in B’-D’, right panel. R^2^ values in left-hand panel of B’-D’ represent the fit for a quadratic polynomial for 72 hours AEL, and straight line for 96 and 120 hours AEL, in the right-hand panel, R^2^ values indicate fit for a quadratic polynomial line for all stages. The fitted curves are plotted with means and 95% confidence intervals. All R^2^ values are presented in Figure S2E. (E) Mitotic nuclear velocity at the developmental stages. Each dot represents single tracked nucleus. Calculated from onset of apical movement and cortical mitotic enrichment, up to most-apical position, generally metaphase. (F-F’’) Exemplary time-lapses of mitotic nuclei moving apically at the developmental stages, in line with panels in (B-D), from wing discs expressing Sqh-GFP and His-RFP. White lines present how tracks were obtained for analysis in A-D. White arrows indicate cortical myosin enrichment associated with directed nuclear movement. White asterisks mark the tracked nuclei when in crowded regions. (//) Represents skipped time steps with little nuclear movement in F’’. The end of metaphase is represented at 00:00 minutes (G) Cross-section of wing disc at 120hours AEL stained with DAPI (blue) and phalloidin (green), and schematic of epithelial topology in wing disc fold regions. Region imaged for apically expanded fold highlighted by dark-red dashed-line box and apically constricted fold highlighted by peach dashed-line box. (H-I) Epithelial morphology measurements for fold regions. (J-J’) Averaged tracking of mitotic nuclei in the fold regions from wing discs expressing Sqh-GFP and His-RFP and obtained as in A. Black dotted line highlights inset in J’, enlarging region with directed apical movement. All tracks are presented in Figure S2F and S2G. (K) Mitotic nuclear velocity for each track in the wing disc folds. (A-E) n = 28, 33 and 34 nuclear trajectories from 3, 3 and 4 wing discs for 72, 96 and 120hr respectively. (H-K) n = 14 and 20 nuclear trajectories from 3 wing discs per condition for AEF and ACF respectively. Scale bars in F are 5*µ*m. Statistical significance in E is given for one-way ANOVA. Statistical significance in I-K is given for unpaired t-test with Welch’s correction for differences in SD. n.s. p > 0.05 ***p < 0.001, ****p < 0.0001. Error bars represent mean ± SEM in A, A’ J, and J’, mean and 95% confidence intervals for B’-D’ and mean ± SD in E, H, I and K.

To quantitatively assess whether the distinct phases in the nuclear trajectories represented distinct modes of nuclear motion we carried out mean square displacement (MSD) analysis on the individual trajectories for each developmental stage. First, we divided the trajectories into “early” and “late” phases as described in the methods (Figures 2B-D). We then calculated the average MSD of the nuclear trajectories in the early and late phases (Figures 2B’,C’,D’). To assess the modes of nuclear motion, we tested fits to the MSD, as a linear MSD verses time lag profile indicates random diffusive movement, a second degree quadratic MSD profile with a positive curvature indicates directed movement, while a negative curvature is indicative of constrained diffusive movement. We found at 72 hours AEL, the early phase MSD profile was better fitted with a second degree quadratic curve with negative curvature, indicating a constrained diffusion of nuclei, in line with the short epithelial structure, short nuclear layer and therefore limited space for nuclear fluctuations (Figure 2B’). For 96 and 120 hours AEL, the early phase MSD to time lag profile fitted a linear function (R^2^ >0.99), indicative of stochastic diffusive motion (Figures 2C’,D’ and S2E). In contrast, during the late phase at all developmental stages, the MSD were best fitted with quadratic curves with positive curvature, indicative of directional movement (Figures 2B’,C’,D’ and S2E). Together, our findings show directed apical nuclear movement is concomitant with mitosis, and therefore might be driven by mitotic rounding.

### Mitotic nuclear dynamics depend on tissue architecture

We next asked whether differences in nuclear dynamics between developmental stages could explain the differences in mitotic positioning observed in fixed tissues (Figures 1E-F’’). First, we compared nuclear velocities specifically during the mitotically-driven, persistent apical movement at each developmental stage. We found that the average mitotic nuclear velocity was greatest at 96 hours AEL (0.89 ± 0.39 *µ*m/min), compared to 72 (0.54 ± 0.21 *µ*m/min) and 120 hours AEL (0.59 ± 0.19 *µ*m/min) (Figure 2E), consistent with observations of instantaneous velocity (Figures 2B,C,D). Together, our results suggest development is associated with differences in nuclear dynamics, in line with observations from our fixed tissue analysis (Figures 1E’,E’’ and S1G-G’’), and may be resultant from changes in tissue architecture.

To further explore how tissue architecture affects nuclear dynamics, we exploited the folds forming in the wing disc at later developmental stages, where the disc locally displays strongly contrasting epithelial morphologies. The fold region possesses distinct folds that have opposing curvature (Figure 2G). At the apical side of the disc, the exposed fold (thereafter apically expanded fold, AEF) has cells with a larger apical than basal surface (Figure 2G). On the basal surface of the disc, cells in the exposed fold (thereafter apically constricted fold, ACF) have a smaller apical than basal surface area (Figure 2G). We imaged the AEF and ACF folds separately in discs at 120 hours AEL, expressing Sqh-GFP and His-RFP. We first quantified the epithelial morphology and revealed differences in nuclear layer heights (Figure 2H), and a greater nuclear density in the AEF compared to the ACF (Figure 2I). We then tracked nuclei within the fold regions and found striking differences in nuclear dynamics (Figures 2J-K and S2F-I). We observed a significantly greater apical mitotic nuclear velocity in the ACF, at 0.77 ± 0.22 *µ*m/min versus 0.35 ± 0.15 *µ*m/min for the AEF (Figure 2K).

Together, our analysis suggests that epithelial morphology can influence mitotic nuclear dynamics.

### Perturbing cell density influences mitotic nuclear positioning

To directly test whether cell density regulates mitotic nuclear behaviour, we genetically and mechanically perturbed cell density and quantified nuclear position in fixed tissues. We first took advantage of a phenotype in the wing disc produced by the reduction of Perlecan (Trol), a component of the extracellular matrix. Expression of *trol-*RNAi has been shown to reduce wing disc surface area, while not affecting the number of cells in the adult fly wing [34, 35]. We characterised how *trol-*RNAi influenced the apico-basal architecture of the tissue at 96 hours AEL and found that the cells redistributed their volume along the apico-basal axis, altering their nuclear layering and tissue packing (Figures 3A-C). Wing discs expressing *trol*-RNAi at 96 hours AEL showed an increased epithelial height (Figures 3B and S3A), nuclear layer height (Figures 3B and S3B) and nuclear density (Figure 3C) compared to age-matched controls. We then assessed the positioning of PH3+ nuclei with *trol*-RNAi and found a significant shift in the distance of mitotic nuclei away from the apical surface compared to age matched controls (Figures 3D and S3C-D’). When the absolute distance of PH3+ nuclei was normalised to differences in the height of the nuclear region, we preserved the shift in mitotic positioning away from the apical surface (Figure S3D,D’). This suggests that the distance mitotic nuclei can migrate is unlikely to give rise to differences in mitotic position, and instead may be due to differences in cell density.

**Figure 3.**
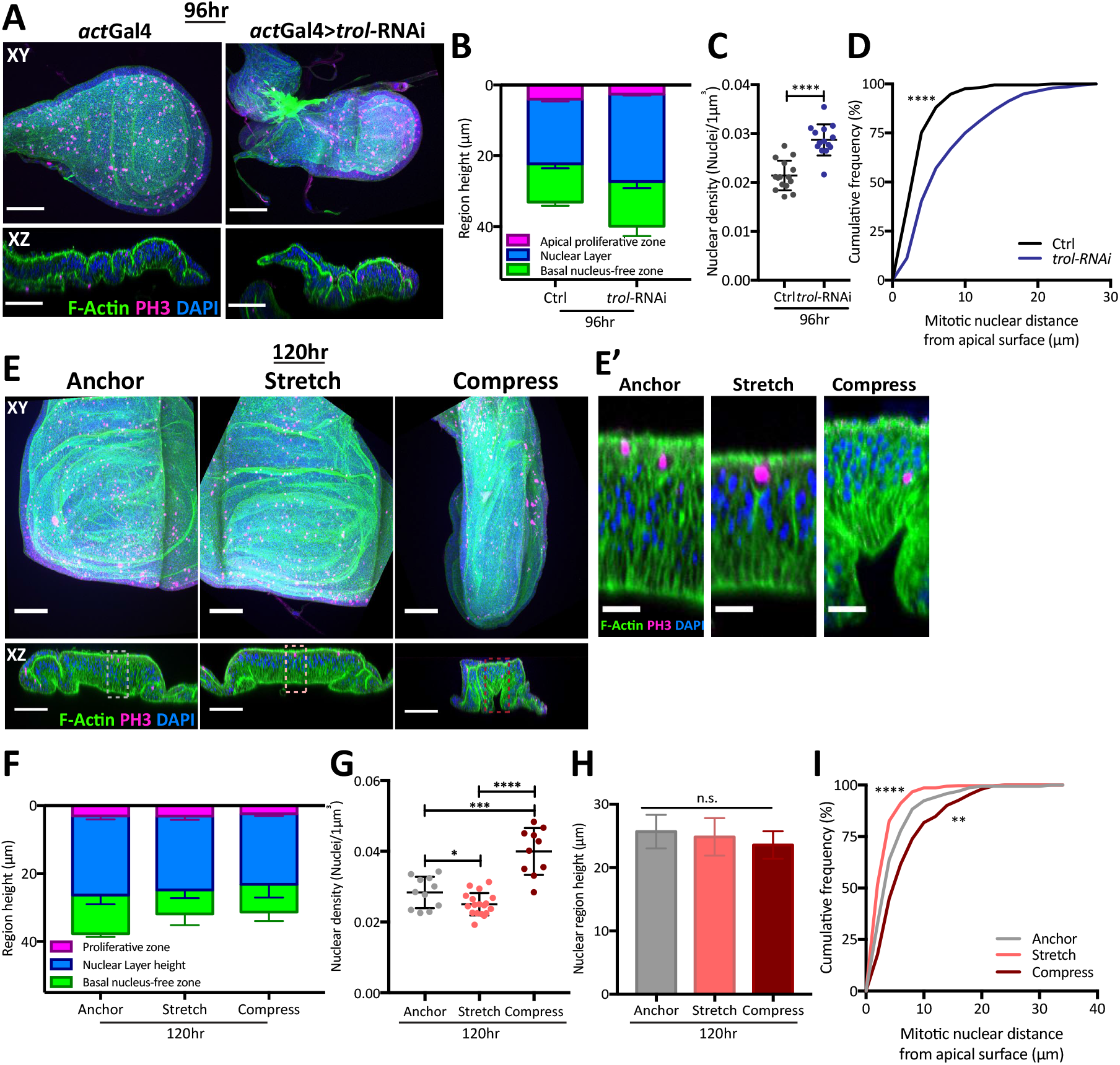
Perturbing cell density influences mitotic nuclear positioning. (A) Wing discs stained with phalloidin (green), anti-PH3 (magenta) and DAPI (blue). The left hand panel presents the wing discs expressing only *actin*Gal4, whilst the right hand panel shows a wing disc expressing *UAS-trol-* RNAi (*perlecan*) under the control of *actin*Gal4. Top view shows projection images, lower images show single lateral cross sections. (B) Quantification of the apico-basal epithelial morphology in the pouch region, for control (*actin*Gal4) and U*AS-trol*-RNAi expressing wing discs. Data individually presented in Figure S3A,B. (C) Nuclear density per *µ*m^3^ for control (*actin*Gal4) and U*AS-trol*-RNAi expressing wing discs. (D) Cumulative frequency distribution of mitotic nuclear distance from the apical surface for control (*actin*Gal4) and U*AS-trol*-RNAi wing discs. Corresponding scatter plots and relative positions presented in Figure S3C-D’. (E) Wildtype 120hr AEL *Drosophila* wing discs, stained with phalloidin (green), anti-PH3 (magenta) and DAPI (blue) and exposed to 30-minute mechanical perturbation using a mechanical tissue-manipulation device. Anchor represents disc within device in a relaxed position. Top view shows projection images, lower images show single lateral cross sections. (E’) Enlarged view of lateral cross sections in E to highlight changes to cell height and nuclear density. (F) Quantification of the apico-basal epithelial morphology in the pouch region of discs upon mechanical perturbation. Data individually presented Figure S3E-F. (G) Nuclear density per *µ*m^3^ for anchored, stretched and compressed conditions in wing disc pouch region. (H) Quantification of nuclear region height showing no effect on the region that can be occupied by nuclei (nuclear region) in wing disc pouch region. (I) Cumulative frequency distribution of mitotic nuclear distance from the apical surface upon mechanical perturbation. Corresponding scatter plots and relative positions presented in Figure S3G-H’. (B and D) n = 6 and 7 wing discs for control and *trol*-RNAi respectively. (C) n = 3 wing discs per condition (F, H and I) n = 3, 8 and 6 wing discs for anchor, stretch and compress conditions respectively. (G) n = 3 wing discs per condition. Scale bars: A,E 50 *µ*m, E’, 10 *µ*m. Statistical significance in C is given for an unpaired t-test, D and I, for Kolmogorov-Smirnov comparison of cumulative distribution and G and H, for one-way ANOVA. *p < 0.05, ***p < 0.001, ****p < 0.0001. Error bars represent mean ± SD in all plots.

To test if cell density was indeed a key feature affecting nuclear dynamics, we developed a manipulation through which we could acutely change cell density, while minimally affecting the nuclear layer height. To this aim, we used a mechanical device [36] to stretch or compress the tissue (Figure 3E), which lead to a reduction and increase in cell density, respectively (Figure 3). Wildtype discs were cultured *ex vivo* within the device for 30 minutes in anchored (unperturbed), stretched or compressed conditions (Figure 3E). We found that whilst the height of the whole tissue was altered when wing discs were stretched or compressed, compared to anchored control wing discs (Figures 3F and S3E), the nuclear region and the nuclear layer were not significantly affected (Figures 3H and S3F). Analysis of PH3+ nuclei position revealed that reducing cell density through tissue stretching, caused a shift in mitotic position towards the apical surface when compared to the anchored control discs (Figures 3I and S3G-H’). Conversely, compressing the tissue caused a marked shift of mitotic nuclei away from the apical surface (Figures 3I and S3G-H’). These results strongly support a direct role for cell density in regulating mitotic position within the tissue and therefore, a likely role in regulating nuclear dynamics.

### Rok is required for apical mitotic nuclear movement at all stages of development

We next asked whether the molecular machinery driving nuclear movements changes as the cell density changes with development. We first explored the role of Rok, a known effector of mitotic nuclear movement in the wing disc [26] and the zebrafish neuroepithelium [24, 25], which is also required to generate force for mitotic rounding in confined cells and tissues [3, 4, 8, 10]. We expressed *Rok-*RNAi using an *engrailed-Gal4* driver in half the wing disc, the other half providing a control for developmental differences in tissue architecture, and assessed mitotic positioning at 72, 96 and 120 hours AEL (Figures 4A and S4A-B’’). We found that in Rok-depleted wing disc epithelia, mitotic nuclei were distributed further from the apical surface compared to the age-matched controls at all developmental stages (Figures 4B and S4C). Furthermore, when mitotic nuclear distances were normalised to the height of the nuclear region, we observed a similar degree of shift in mitotic positioning at each stage, suggesting that Rok is equally important at all three developmental stages investigated (Figures 4B’ and S4C’).

**Figure 4.**
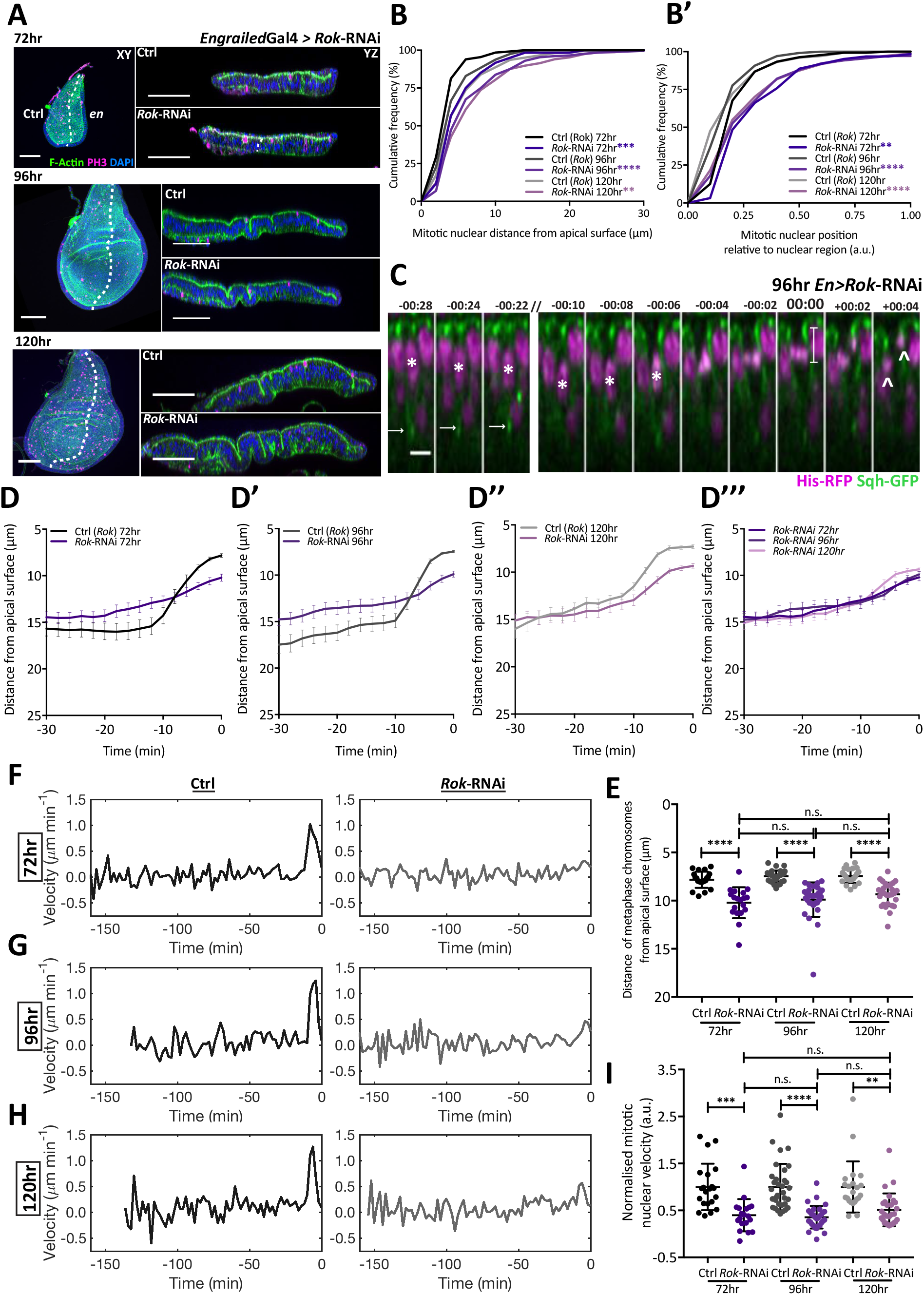
Rok is required for apical nuclear movement at all stages of development. (A) Representative images of wing disc morphology with *Rok*-RNAi in the engrailed domain (posterior, en) at different developmental stages and stained with phalloidin (green), anti-PH3 (magenta) and DAPI (blue). Left images represent projection images, right images show lateral cross sections along the dorsal-ventral axis (white dashed lines show anterior-posterior boundary), from the centre of anterior half and the posterior half. (B) Cumulative frequency distribution of mitotic nuclear distance from apical surface for control (anterior) and *Rok*-RNAi (posterior) wing disc regions at different developmental stages. (B’) Cumulative frequency distribution of mitotic nuclear position, relative to the nuclear region, for anterior control and posterior *Rok*-RNAi expressing wing disc regions at different developmental stages. (B-B’) Data corresponds to Figure S4C-C’. (C) Representative time-lapse of a mitotic nucleus moving apically within *Rok*-RNAi expressing, posterior wing pouch region, at 96hours AEL in a wing disc co-expressing Sqh-GFP and His-RFP. White arrows indicate cortical myosin enrichment. White asterisks mark the tracked nucleus when in crowded regions. (//) Represents skipped time steps with little nuclear movement. 0 minutes marks the timepoint immediately prior to anaphase, the white line shows the final distance of the metaphase chromosomes from the apical surface, for which values are presented in E. The last two timepoints illustrate failure to orient planar cell division (arrowheads highlight separating chromosomes). (D-D’’’) Average nuclear tracks for final 30 minutes prior to final metaphase, for anterior control and posterior *Rok*-RNAi mitotic cells, at different developmental stages. Full time course shown in Figure S4F-G. (E) The final distance of the metaphase chromosomes from the apical surface for control and *Rok*-RNAi expressing mitotic cells and at each developmental stage. Measurement obtained from the position at 0 minutes as illustrated in C. (F-H) Average instantaneous velocity measurements for tracked nuclei in anterior control and posterior *Rok*-RNAi mitotic cells and at each developmental stage, from full time course shown in Figure S4F-G. (I) Mitotic nuclear velocity for control and *Rok*-RNAi expressing mitotic cells, normalised to average rate of apical nuclear movement for age-matched controls. Each dot represents single tracked nucleus. Calculated from onset of apical movement and cortical mitotic enrichment, up to most-apical position, i.e. metaphase. (B-B’) n = 8, 7 and 5 wing discs for 72, 96 and 120 hours respectively (D-I) n = 19-33 nuclear trajectories acquired from 3 wing discs per developmental stage. Scale bars: A, 50 *µ*m, C, 5 *µ*m. Statistical significance given for Kolmogorov-Smirnov comparison of cumulative distribution in B-B’ and one-way ANOVA in E-I. n.s. p > 0.05, *p < 0.05, **p < 0.01, ***p < 0.001, ****p < 0.0001. Error bars represent mean ± SEM in D-D’’’ and mean ± SD in E and I.

To gain further insight into the role of Rok in regulating apical mitotic nuclear movement, we assessed how the nuclear dynamics were perturbed during development using live imaging. As for wildtype analysis, we tracked nuclei within the pouch region, expressing Sqh-GFP and His-RFP (Figure 2), in addition to *Rok*-RNAi in the *engrailed* compartment (Figures 4A,C and S4D,E). We observed a consistent defect in nuclear trajectories with *Rok*-RNAi compared to age-matched controls, but little difference between the developmental stages (Figures 4C-D’’’ and S4D-G). To assess the success of apical mitotic migration, we plotted the final distance of the metaphase chromosomes from the apical surface. We found a significant defect in apical positioning with *Rok*-RNAi compared to controls at all developmental stages (Figure 4E). Subsequently, cells with non-apical mitotic nuclei often divided along the apico-basal axis, rather than being planar-orientated (Figures 4C, S4D,E, white arrow heads), supporting previous work investigating the importance of Rok and mitotic rounding for planar-oriented division [9, 10]. We also observed a reduced apical mitotic nuclear velocity compared to internal controls (Figure S4H), consistent with the plots of instantaneous nuclear velocity (Figures 4F-H). We normalised the mitotic nuclear velocity by the average apical mitotic nuclear velocity of the control region and found a comparable relative reduction in the normalised velocities with *Rok*-RNAi at each developmental stage (Figure 4I). Although expressing *Rok*-RNAi appeared to globally delay wing disc development compared to wildtype tissue, we confirmed that nuclear density was increasing incrementally at each developmental stage in the *Rok*-RNAi compartment (Figures 1D and S4I).

Together, these results show that Rok is essential for efficient apical mitotic nuclear movement at all developmental stages examined, suggesting that the requirement for Rok is not influenced by differences in epithelial architecture.

### Dependency on Dia for apical mitotic nuclear movement increases with development

Actin destabilising drugs have been shown to perturb apical mitotic positioning in the wing disc and the zebrafish neuroepithelium [24–26]. Interestingly, formin-mediated actin polymerisation is also essential for mitotic rounding when cells are under confinement [2–4, 8]. We therefore investigated the role of Dia, a *Drosophila* formin, in regulating apical mitotic positioning as cell density increases with development. As for *Rok*-RNAi, we reduced Dia expression in half the wing disc by expressing *dia*-RNAi under the control of the *engrailed* driver, maintaining an unperturbed internal control for comparison (Figures 5A and S5A-B’’). We first assessed the effects on mitotic positioning in fixed tissue. Upon *dia*-RNAi, we found mitotic nuclei were distributed further from the apical surface at 96 and 120 hours AEL, than at 72 hours AEL, compared to their age-matched controls (Figures 5B,B’ and S5C,C’). These observations suggest that there might be development specific differences in the requirement for Dia in nuclear dynamics.

**Figure 5.**
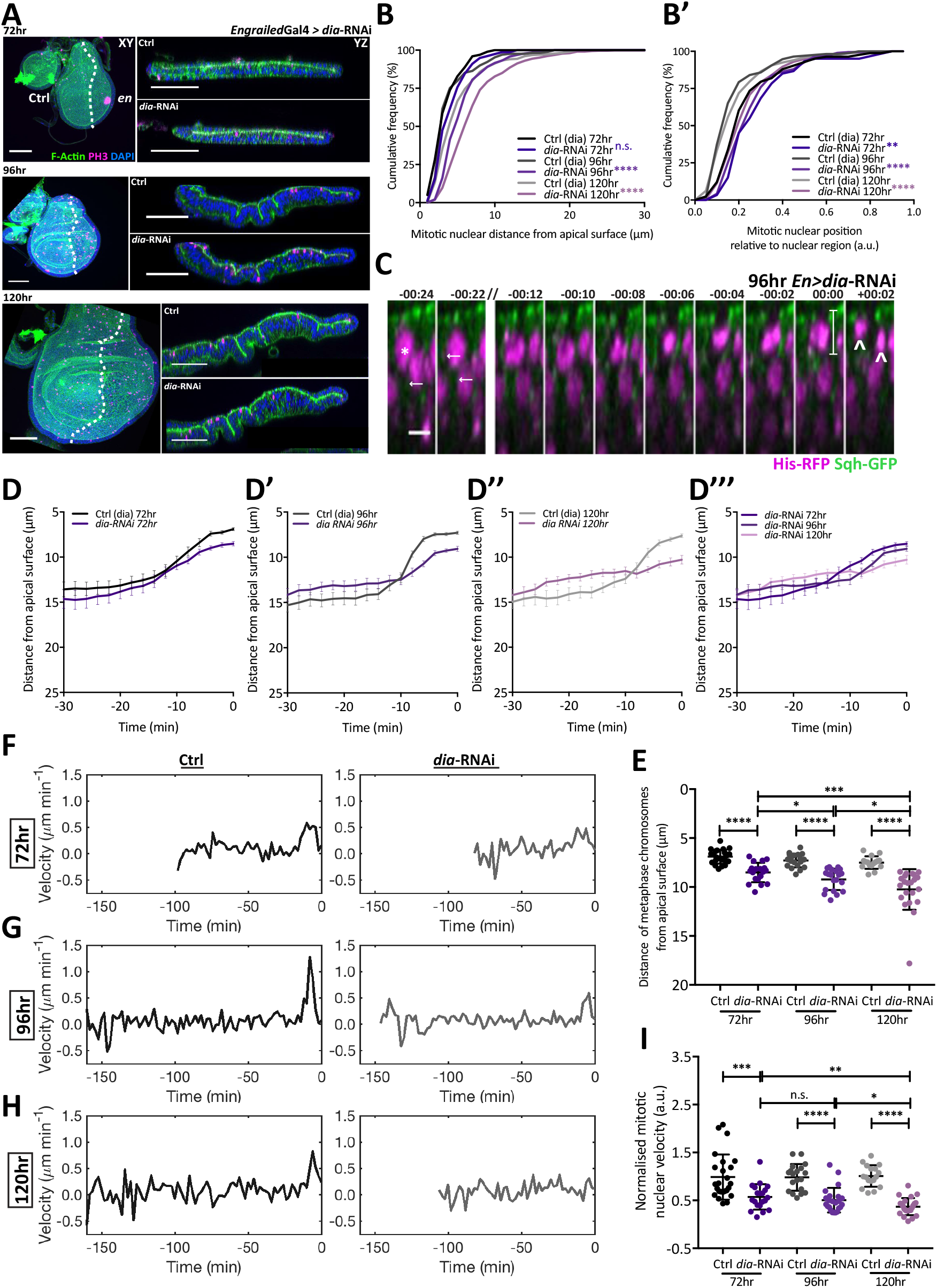
Dependency on Dia for apical mitotic nuclear movement increases with development. (A) Representative images of wing disc morphology with *dia*-RNAi in the *engrailed* domain (posterior, *en*) at different developmental stages and stained with phalloidin (green), anti-PH3 (magenta) and DAPI (blue). Left images represent projection images, right images show lateral cross sections along the dorsal-ventral axis (white dashed lines show anterior-posterior boundary), from the centre of anterior half and the posterior half. (B) Cumulative frequency distribution of mitotic nuclear distance from apical surface for control (anterior) and *dia*-RNAi (posterior) wing disc regions at different developmental stages. (B’) Cumulative frequency distribution of mitotic nuclear position, relative to the nuclear region, for anterior control and posterior *dia*-RNAi expressing wing disc regions at different developmental stages. (B-B’) Data corresponds to Figure S5C-C’. (C) Representative time-lapse of a mitotic nucleus moving apically within *dia*-RNAi expressing, posterior wing pouch region, at 96hours AEL in a wing disc co-expressing Sqh-GFP and His-RFP. White arrows indicate cortical myosin enrichment. White asterisks mark the tracked nucleus when in crowded regions. (//) Represents skipped time steps with little nuclear movement. 0 minutes marks the timepoint immediately prior to anaphase, the white line shows the final distance of the metaphase chromosomes from the apical surface, for which values are presented in E. The last two timepoints illustrate failure to orient planar cell division (arrowheads highlight separating chromosomes). (D-D’’’) Average nuclear tracks for final 30 minutes prior to final metaphase, for anterior control and posterior *dia-*RNAi mitotic cells, at different developmental stages. Full time course shown in Figure S5F-G. (E) The final distance of the metaphase chromosomes from the apical surface for control and *dia*-RNAi expressing mitotic cells and at each developmental stage. Measurement obtained from the position at 0 minutes as illustrated in C. (F-H) Average instantaneous velocity measurements for tracked nuclei in anterior control and posterior *dia-*RNAi mitotic cells and at each developmental stage, from full time course shown in Figure S5F-G. (I) Mitotic nuclear velocity for control and *dia*-RNAi expressing mitotic cells, normalised to average rate of apical nuclear movement for age-matched controls. Each dot represents single tracked nucleus. Calculated from onset of apical movement and cortical mitotic enrichment, up to most-apical position, i.e. metaphase. (B-B’) n = 4, 7 and 8 wing discs for 72, 96 and 120 hours respectively (D-I) n = 16-24 nuclear trajectories acquired from 3-4 wing discs per developmental stage. Scale bars: A, 50*µ*m, C, 5*µ*m. Statistical significance given for Kolmogorov-Smirnov comparison of cumulative distribution in B-B’ and one-way ANOVA in E-I. n.s. p > 0.05, *p < 0.05, **p < 0.01, ***p < 0.001, ****p < 0.0001. Error bars represent mean ± SEM in D-D’’’ and mean ± SD in E and I.

We then tracked nuclei in *dia*-RNAi expressing wing discs and analysed the nuclear dynamics of mitotic cells. We observed a significant defect in mitotic nuclear dynamics upon Dia depletion at 72, 96 and 120 hours AEL, however, visual inspection of the trajectories suggested the defects increased through development (Figures 5C-D’’’ and S5D-G). To explore this further, we first assessed the success of apical positioning by plotting the final distance of the metaphase chromosomes from the apical surface. We found the distance from the apical surface at metaphase was significantly higher in dia-depleted wing discs compared to age-matched controls, and this defect increased through development (Figure 5E). We then analysed whether the nuclear velocity at mitosis was affected by Dia depletion. We observed a reduced mitotic nuclear velocity (Figure S5H) and instantaneous velocity at the late phase (Figures 5F-H) in Dia-depleted epithelia, compared to internal controls. To assess whether there were differences in the extent of the effect between developmental stages, we normalised the mitotic nuclear velocities to the average mitotic nuclear velocity of the age-matched control region. We found that the difference between the normalised velocities with *dia-*RNAi increased during development, with the greatest reduction in normalised velocity at 120 hours AEL (Figure 5I) when nuclei are most tightly packed (Figure S5I). As in Rok-depleted wing discs, when nuclei failed to reach the apical surface due to *dia*-RNAi expression, they often exhibited defects in planar-oriented cell division (Figures 4C and S4E, white arrow heads).

It has previously been shown that Arp2/3-mediated actin polymerisation is not required for mitotic rounding upon confinement [3]. We therefore asked whether Arp2/3 is also dispensable for mitotic nuclear movement. We used the Arp 2/3 complex inhibitor CK666 to treat 120-hour discs expressing the cell-membrane marker Basigin-GFP, and His-RFP to mark nuclei, and tracked nuclear movement. We found that inhibiting Arp2/3 actin polymerisation had no significant effect on nuclear dynamics (Figures S5J-L).

Together, our results suggest that apical mitotic nuclear migration increasingly depends on Dia-mediated actin polymerisation as development progresses.

### Defects in nuclear positioning upon Dia depletion can be rescued by mechanically reducing cell density

We subsequently asked whether the increasing cell density in the developing epithelium was responsible for the worsening defect in mitotic nuclear dynamics upon Dia depletion. To address the role of cell density in dependency on Dia, we investigated whether decreasing cell density in *dia*-RNAi discs would rescue defects in mitotic nuclear movement. We utilised our mechanical device to reduce cell density through tissue stretching and assessed mitotic nuclear positioning (Figures 6A,B and S6A,B). Wing discs at 120 hours AEL expressing *dia*-RNAi in whole wing pouch region were stretched for 30 minutes and then fixed. We then measured mitotic nuclear positions and observed a significant shift in the distribution of mitotic nuclei towards the apical surface upon mechanical stretch in both control and discs expressing *dia*-RNAi (Figures 6C-C’’ and S6C-C’’). This suggests that mechanically reducing cell density can partially rescue defective apical mitotic positioning produced by Dia depletion.

**Figure 6.**
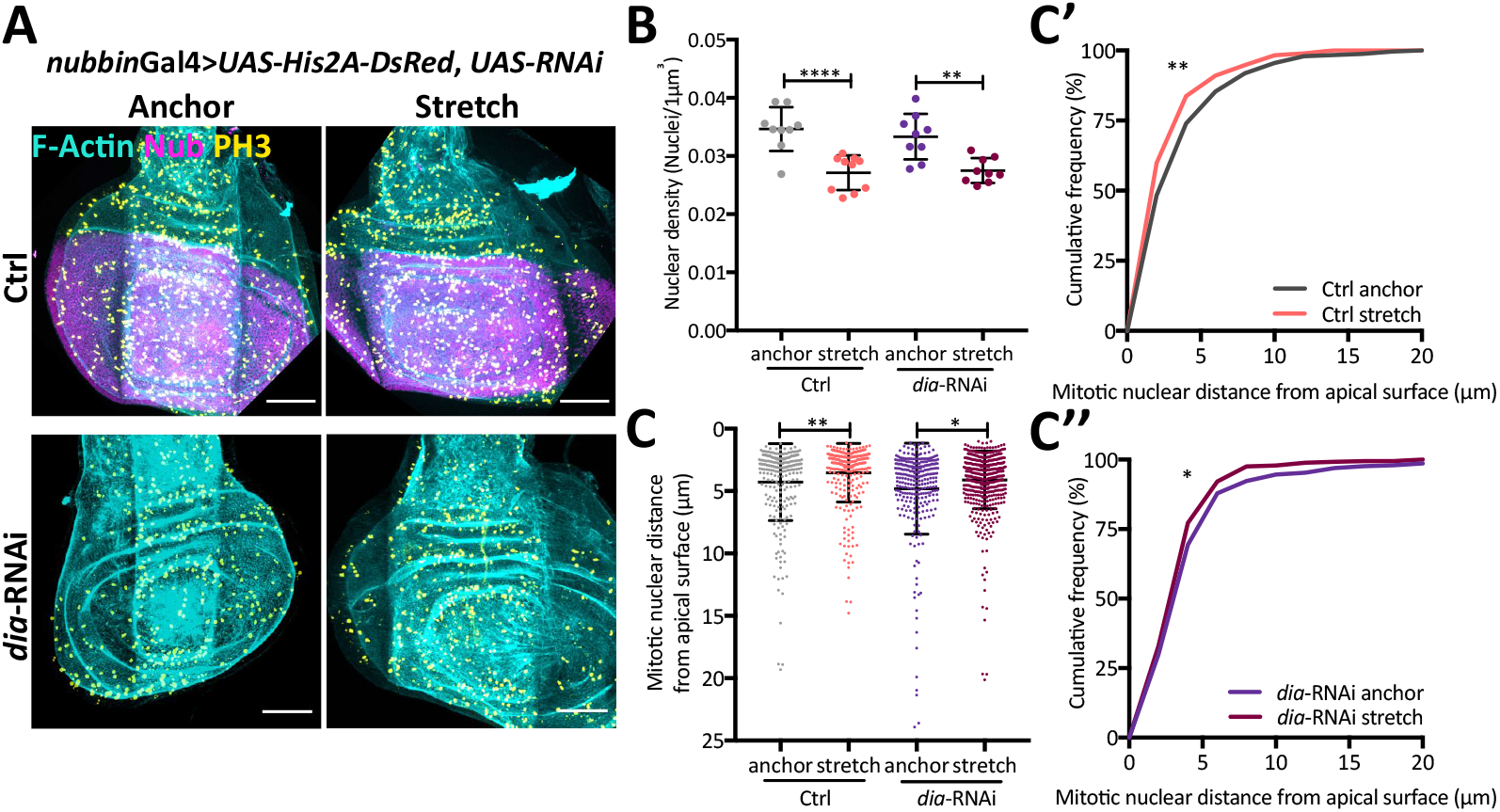
Defects in nuclear positioning upon Dia depletion can be rescued by mechanically reducing cell density. (A) Representative images of mechanical stretch with control and *dia*-RNAi expression under the control of a *nubbin*-Gal4 driver, co-expressing UAS-His2A (magenta), stained with phalloidin (cyan) and anti-PH3 (yellow). (B) Density of nuclei per 1*µ*m^3^ in the pouch region for control and *dia*-RNAi expressing discs in anchor and stretched conditions. (C) Mitotic nuclear distance from apical surface in the pouch region of the wing disc for control and *dia*-RNAi expressing wing discs in anchor and stretched conditions, presented as a dot plot. (C’-C’’) Data in C presented as a cumulative frequency distribution. Normalised data of C-C’’ presented in Figure S6C-C’’. (B) n= 3 wing discs per condition (C-C’’) n = 6 and 6 wing discs for control anchor and stretch respectively and n = 7 wing discs each for *dia*-RNAi anchor and stretch. Scale bars: A, 50*µ*m. Statistical significance given for Kolmogorov-Smirnov comparison of cumulative distribution. *p < 0.05, **p < 0.01, ****p < 0.0001. Error bars represent mean ± SD in B and C.

Together, our results show that in wing discs expressing *dia*-RNAi, the increasing defect in apical mitotic positioning and nuclear velocity as development progresses is likely due to the increasing cell density.

## Discussion

Our findings demonstrate that nuclear movement required for apical mitosis in the pseudostratified *Drosophila* wing disc is mechanically regulated by tissue architecture. We use live imaging of the developing wing disc, in combination with analysis of fixed samples where the cell cycle stage can be unambiguously determined. We confirm through the combinatorial use of live and fixed analysis that in the wing disc, nuclear movement in mitosis, rather than G2 [25, 28, 29], achieves apical mitotic positioning [26] (Figure 2). Furthermore, our fixed-tissue analysis of mitotic nuclear position recapitulates developmental stage specific differences in mitotic nuclear dynamics (Figures 1E’-E’’, 2 and S1G’-G’’), thereby allowing us to assess mitotic nuclear behaviours using fixed samples when mechanical and genetic perturbations of tissue architecture were not amenable to live imaging (Figure 3).

We show that mitotic nuclear dynamics change as development progresses and tissue architecture changes (Figures 1 and 2). Previous studies had suggested that tissue density might influence IKNM in the crowded, vertebrate cortex PSE [28, 29]. Here, we used genetic and mechanical perturbations to acutely modify tissue packing and directly show that nuclear density affects mitotic nuclear positioning (Figure 3). Our data thus strongly suggests that the changes in nuclear density that accompany development mediate differences in nuclear dynamics.

Interestingly, we observe that apical mitotic nuclear velocity is lower in 72-hour wing discs, compared to 96-hour wing discs, even though pseudostratification and nuclear packing are lower at 72 hours AEL. The wing disc at 72 hours AEL is not yet clearly pseudostratified and the cells display simple, columnar-like morphology. It is possible that due to this morphology, mitotic rounding, which requires the cells to bring the lateral membranes together, is overall slower than at 96 hours AEL, when pseudostratification brings the cell membranes in close proximity even in non-mitotic cells (Figure 2F-F’’).

Notably, we also show that tissue folds dramatically alter nuclear dynamics, likely due to resultant differences in nuclear density. The positive apical curvature of the wing disc AEF increases nuclear crowding at the apical surface, leading to a reduced mitotic nuclear velocity compared to the ACF, in which the negative apical curvature displaces nuclei basally. Tissue curvature and folding exists in many PSE, including the developing intestine [12], the zebrafish retina [37] and the nervous system. It will be interesting to investigate whether curvature similarly affects nuclear organisation and IKNM dynamics in other tissues.

Our observations suggest a mechanism by which mechanically regulated nuclear dynamics as tissue-packing increases could impact tissue growth. Previous studies of PSE in the developing nervous system have hypothesised that tissue packing and proliferation are linked [13, 33], but no mechanism has so far been identified. Interestingly, at 120 hours AEL when we observe a significantly slowed mitotic nuclear velocity (Figure 2), the wing disc is approaching growth arrest. Previous studies of wing disc growth demonstrate a marked increase in cell cycle length and a reduced mitotic index at 120 hours AEL compared to wing discs at 72-96 hours AEL [32, 38]. Our findings indicate that increasing cell density could increase the length of mitosis, as apical migration is delayed, suggesting a possible mechanism directly linking tissue packing and cell cycle length.

Finally, we show that reducing Rok and Dia in the wing disc leads to reduced mitotic nuclear velocity, mispositioning of mitotic nuclei away from the apical surface and a greater frequency of sub-apical divisions (Figures 4 and 5). The requirement for Rok, a key activator of Myosin II (myosin), does not appear to depend on developmental stage, suggesting that increased myosin activity is required for apical mitotic migration independently of the level of tissue packing. This is in line with a general requirement for myosin activity to support mitotic rounding [4, 39, 40]. Consistently, work in shorter and less densely packed epithelia has shown that Rok depletion leads to defects in mitotic rounding and, as a result, division defects [3, 8, 10].

In contrast, the effects of Dia depletion worsen at later stages of development. Our data suggests that the increased requirement for Dia as development progresses is linked to the concomitant increased cell density, since mechanically reducing density partially rescues mitotic nuclei positioning defects in Dia depleted discs (Figure 6). Interestingly, Dia has been shown to be required for effective tension generation in the actomyosin cortex [5] and is particularly important for cell rounding in confinement [2–4]. The increased dependency on Dia for apical nuclear migration therefore suggests that nuclear movement and mitotic rounding are likely coupled, supporting our live analysis of persistent apical motion upon mitosis. Furthermore, these results suggest that with greater cell density, there is a greater force requirement for apical mitotic rounding in PSE.

Interestingly, genetically increasing Rok activity in the wing disc has been shown to increase cell numbers, while reducing Rok activity reduces cell numbers [41]. Therefore, regulating mitotic nuclear dynamics could be tightly linked to wing disc growth control. How Dia expression influences cell numbers in the wing disc is open to further research.

In summary, our results reveal how the mechanical environment imposed on cells confined within a tissue can influence molecular and cellular mechanisms regulating nuclear movement. Future studies will be required to dissect how exactly feedback between tissue architecture and single cell behaviour controls global tissue growth and morphogenesis.

## Acknowledgements

We thank Robert Tetley for critical comments on the manuscript. We thank the Sanson, Baum and Thompson labs for fly lines and the LMCB core microscopy facility for imaging support. NJK is funded by a MRC PhD studentship. MT is funded by a Sir Henry Wellcome Fellowship (Grant No: 103095). YM is funded by a MRC Fellowship MR/L009056/1, a UCL Excellence Fellowship, a NSFC International Young Scientist Fellowship 31650110472 and a Lister Institute Research Prize Fellowship. This work was also supported by MRC funding to the MRC LMCB University Unit at UCL, award code MC_U12266B.

## Author Contributions

Author contributions: NJK designed the research, performed the experiments, analysed the data and wrote the manuscript. ACY helped with immunostaining experiments. MT performed velocity, mean square displacement analysis and wrote the manuscript. NH helped to obtain fold region nuclear trajectories. EP and YM designed the research and wrote the manuscript.

## Materials and Methods

### Contact for reagent and resource sharing

Further information and requests for resources and reagents should be directed to and will be fulfilled by Dr. Yanlan Mao and Prof Ewa Paluch.

### Experimental model and subject details

#### Drosophila melanogaster

Fly stocks were raised in non-crowded conditions on standard cornmeal molasses fly food medium at 25°C. Fly food consisted of, per 1L, 10g agar, 15g sucrose, 33g glucose, 35g years, 15g maize meal, 10g wheat germ, 30g treacle, 7.22g soya flour, 1g nipagin, 5ml propionic acid.

Male and female larvae were dissected at early to late 3rd instar development (approximately 72-120hr AEL) for experiments. For developmental staging, flies were staged every 24 hours. Appropriate larva were selected for dissection and wing disc morphology used to refine staging. 72-hour discs were defined by flat, tear drop shaped epithelia; 96-hour wing discs had all three folds formed but their surface area measures substantially smaller than the later stages (30,000*µ*m2); 120-hour wing discs have all three folds, are enlarged (46,000*µ*m2) and exhibit a condensation of cells at the dorsal-ventral midline. For trol-RNAi mutants, in which disc morphology differs, discs were staged to the timing of the wild-type 96-hour counterparts.

Strains used:

For wildtype, fixed experiments presented in Figure 1, S1, 3E-I and S3E-H’: *yw;;;* (Bloomington)

Live imaging of cell shape and nuclear tracking in Figure 2, S2, 4C-I, S4D-H, 5C-I and S5D-H: *;;SqhGFP, HisRFP;* (self-generated)

For expression of *trol*-RNAi in Figure 3A-D and S3A-D’: ;*actGal4/CyO;;* (Baum Lab*), *trol*-RNAi (VDRC: GD/24549)

Perturbation of Rok and Dia expression in Figure 4, S4, 5, S5, 6 and S6:

;EnGal4, ECad-GFP; UAS-NLS-Cherry; (Self-generated with *enGal4* from Sanson Lab), *Rok*-RNAi (VDRC KK/104675), *dia*-RNAi (VDRC KK/103914), ;*enGal4; SqhGFP, HisRFP;* (self-generated using *enGal4* from Sanson Lab), *nubGal4; UAS-His2A-DsRed* (Thompson Lab)

CK666 inhibition of Arp2/3, live imaging in Figure S5I-K: ;Bsg-GFP, His-RFP; (DGRC 115366)

### Method details

#### Immunofluorescence

Larva and forceps were washed in 70% ethanol and PBS. Larva were transferred to dissecting medium (Shields and Sang M3 media, 2% FBS, 1% pen/strep, 3 ng/ml hydroxyecdysone and 0.05 units/l insulin) and their wing discs extracted for up to 15 minutes. Discs were fixed for 30 minutes in 4% formaldehyde-PBS at room temperature. Fixed tissue was washed 4×10 min PBT (PBS, 0.3% Triton X-100) and 4×10 min PBT-BSA (PBT, 0.5% BSA). Primary antibody at appropriate concentrations was incubated overnight. Washed were repeated and secondary antibody, with DAPI and Phalloidin incubated for 1-2 hours at room temperature. Discs were washed for 3×20 min PBT and 3x quick rinse with PBS. Discs were mounted in Fluoromount G slide mounting medium (Southern Biotech) for imaging.

Antibodies and Dyes: Mouse anti-PH3 (Millipore 3H10 05-806), 1:500. Rabbit anti-PH3 (Millipore 3H10 06-570), 1:150. Mouse anti-Engrailed (DSHB, 4D9), 1:50. Goat anti-mouse Alexa-488 (Life Technologies, A11029), 1:500. Donkey anti-mouse-RRX (JacksonImmunoResearch), 1:500. Goat anti-rabbit Alexa-488 (Life Technologies, A11034), 1:500. Goat anti-rabbit Alexa-555 (Abcam, ab150086), 1:500. DAPI (Sigma-Aldrich, D8417), 1:1000. Alexa Fluor 647 Phalloidin (Cell Signalling, 8953S and A22287, Life Technologies) (1:20).

Imaging: Samples were imaged using a Leica SP5 or SP8 inverted confocal microscope with a 40X objective, 1-2X zoom, 0.35 *µ*m depth resolution and 1024^2^ or 512^2^ XY pixel resolution.

#### Live imaging

Pouch region sample preparation: Discs were dissected as described above. Discs were transferred in dissecting media to Fluoro glass-bottomed dish and positioned apical side down onto a 0.4 *µ*l line of Cell-Tak (Corning, 354240) (previously dried onto glass-bottom on heat plate set to 29°C). A further 1 ml of dissecting media was added before sealing the dish with parafilm.

Fold region sample preparation: Discs mounted as described for pouch region with the exception of the basal fold, which is instead mounted basal side down.

CK666 sample preparation: Dissected discs were incubated in 80 *µ*M in dissecting media for 30 minutes prior to assembling for live imaging, as conducted for pouch region imaging.

Imaging: All live imaging was carried out using an inverted Zeiss LSM880 microscope with 40X oil lens, 2X zoom, 512by256 XY resolution and 0.5 *µ*m depth resolution for confocal stack imaging. The LSM880 Airyscan detector was used in confocal setting for sensitive imaging. Laser settings and resolution were identical for each individual experimental group and for *en*Gal4, RNAi expression experiments, both tissue regions were in the same field of view.

#### Wing disc stretch and compression

Larva were dissected as for immunofluorescence and live imaging experiments. The assembly and use of the stretcher device has been previously described[36]. Specific to this work, discs were loaded into the device so the manipulation was applied bi-directionally along the anterior-posterior axis and held for 30 minutes. Compression was applied by pre-stretching PDMS membranes prior to disc loading. The loaded PDMS was then relaxed to compress the tissue along the anterior-posterior axis. Tissues were fixed directly within the device with 4% PFA in PBS, for 15 minutes before transferring discs to a glass-well dish for a further 15 minutes of fixing. Immunostaining is carried out as described above.

### Quantification and statistical analysis

#### Quantification of epithelial morphology

To measure the height of the epithelia, nuclear layer, nuclear region, apical proliferative zone and the basal nucleus free zone, confocal stack images were “resliced” in ImageJ along the dorsal-ventral axis. For yw, mechanically-perturbed and *trol*-RNAi discs, measurements were taken from the anterior third, centre and posterior third of the pouch region. Three intensity profile measurements from the apico-basal axis were obtained from the DAPI channel as shown in Figure S1A, left panel. For mutant perturbation with *EnGal4, ECad-GFP; UAS-NLS-Cherry* three measurements from two cross-sections in the anterior and posterior compartment were acquired, representing the control and mutant respectively. Each respective measurement was obtained by the measurements for the epithelial height, the starting and the final position of the nuclear layer as illustrated in Figure S1A, right panel. The number of nuclei within the nuclear layers was measured by the counting the number of peaks for each intensity profile. The average value of the regions was calculated for each disc and subject to an independent t-test and presented as a plot. The values for the apical proliferative zone, the nuclear layer and the basal nucleus free zone were presented as “Stacked Bars” in Prism to reflect the epithelium morphology for the disc condition (For example see Figure 1B and 1B’).

#### Quantification of nuclear density

Density was calculated as in Bittig et al., 2009 [31]. 3-5 segments of 50-90μm^2^ apical surface areas were sampled per disc and “resliced” to present the z-sectioning in Image J. The volume of the nuclear region was calculated by multiplying the apical surface area measurement by the height of the nuclear layers. 3 wing discs were measured per condition, and each density measurement collated and presented to demonstrate the range across the wing disc.

#### Quantification of mitotic positioning in fixed tissue

In the pouch region of the wing disc, the mitotic distance was obtained by measuring the distance from the centre of the PH3+ nuclei to the apical surface, as marked by F-Actin as illustrated in Figure 1E. To obtain a position with respect to the region in which the mitotic nuclei could translocate, the mitotic distance was divided by the average height of the nuclear region as illustrated in Figure S1G. To plot the distribution of nuclei for each measurement type, the data points from each age-matched disc of an experimental group were pooled. The distribution of the mitotic positions were statistically compared using the non-parametric Kolmogorov-Smirnov comparison of cumulative distribution. For scatter plots, the Y axis scale was reversed so the PH3 position would represent the apico-basal organisation of the wing disc cells as depicted in the images of the epithelium.

#### Nuclear tracking

Confocal stack, time-lapse images were corrected for drift using the 3D drift correction plugin in ImageJ. The corrected imaging was the “resliced” to present the apico-basal axis. Nuclei were manually tracked by measuring the distance from the basal side of the nucleus to the apical surface at each time-step, with nuclei marked by His-RFP and the apical surface marked by Sqh-GFP expression. Mitotic cells were identified by tracking backwards from when the cell is in a final metaphase state, prior to anaphase chromosome separation. Excel and Prism were used to revers and align tracks and present trajectories. Tracks were statistically analysed as described below.

For supplementary movies and montage images the drift corrected time-lapse imaging were subject to background subtraction (rolling ball radius 30, ImageJ) and 3D Gaussian filtering (1.0 pixel diameter) before presenting.

#### Apical mitotic rate determination

The start of apical mitotic movement was determined by the initiation of persistent apical movement and cortical Sqh-GFP enrichment, whilst the final position was determined at the metaphase, assessed from the nuclear tracks and the respective live imaging. The distance between these two points was divided by the length of time taken to calculate the apical mitotic rate for each track.

To compare the fold effect of *dia* and *Rok* RNAi expression on apical mitotic rate, each apical mitotic rate measurement was divided by the average mitotic rate measurement for the control condition at the respective disc age (the average of all tracks, across all discs at the respective developmental stage).

#### Calculation and assessment of mean squared displacement profiles

The mean squared distance of a trajectory for a given time lag Δ ***t*** is calculated from the average of squared displacement all data couples separated by Δ ***t*** For a trajectory of N data points and a data collection interval of δ***t***, the MSD for lag Δ ***t*** = ***n* δ *t*** becomes:

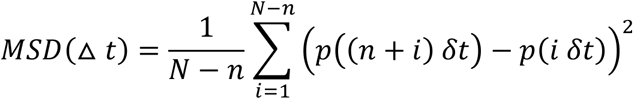

The MSD profiles for increasing time lag were calculated for each individual nucleus sub-trajectory towards the apical surface. The sub-trajectories are defined as the early and late phase nuclear migration. The transition from the early to the late phase was determined manually as the onset of persistent apical movement and enrichment of cortical Sqh-GFP. The late phase was truncated at metaphase when no further apical movement occurred. The maximum lag is limited by the length of the late phase trajectory in each developmental time and mutant data set.

With a power law model 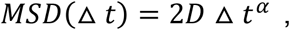 stochastic diffusive motion corresponds to values of α close to 1, making the best fit a linear fit. A directional diffusion with flow model on the other hand fits best with a quadratic model, 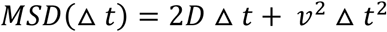.

Linear and second-degree polynomial quadratic fits are calculated for each of early and late phases, with a forced interception at the origin. The goodness of fit is assessed by the R^2^ value of the fitted model, coupled by visual inspection.

#### Statistical analysis

Microsoft Excel 2011, Matlab 2016 and Prism 7 Software were used to present data and conduct statistical analysis. The respective statistical tests are described in the figure legends.

For epithelial morphology analysis, a minimum of three discs were used per condition. For fixed tissue analysis of mitotic position, a minimum of 5 discs were used per condition. For live nuclear tracking, a minimum of three discs were used per condition. The following statistical significance cut off was applied: n.s. p>0.05, * p<0.05, **p<0.01, ***p<0.01, ****p<0.0001.

## Supplementary Information

**Figure S1, Supplementary data for Figure 1:**
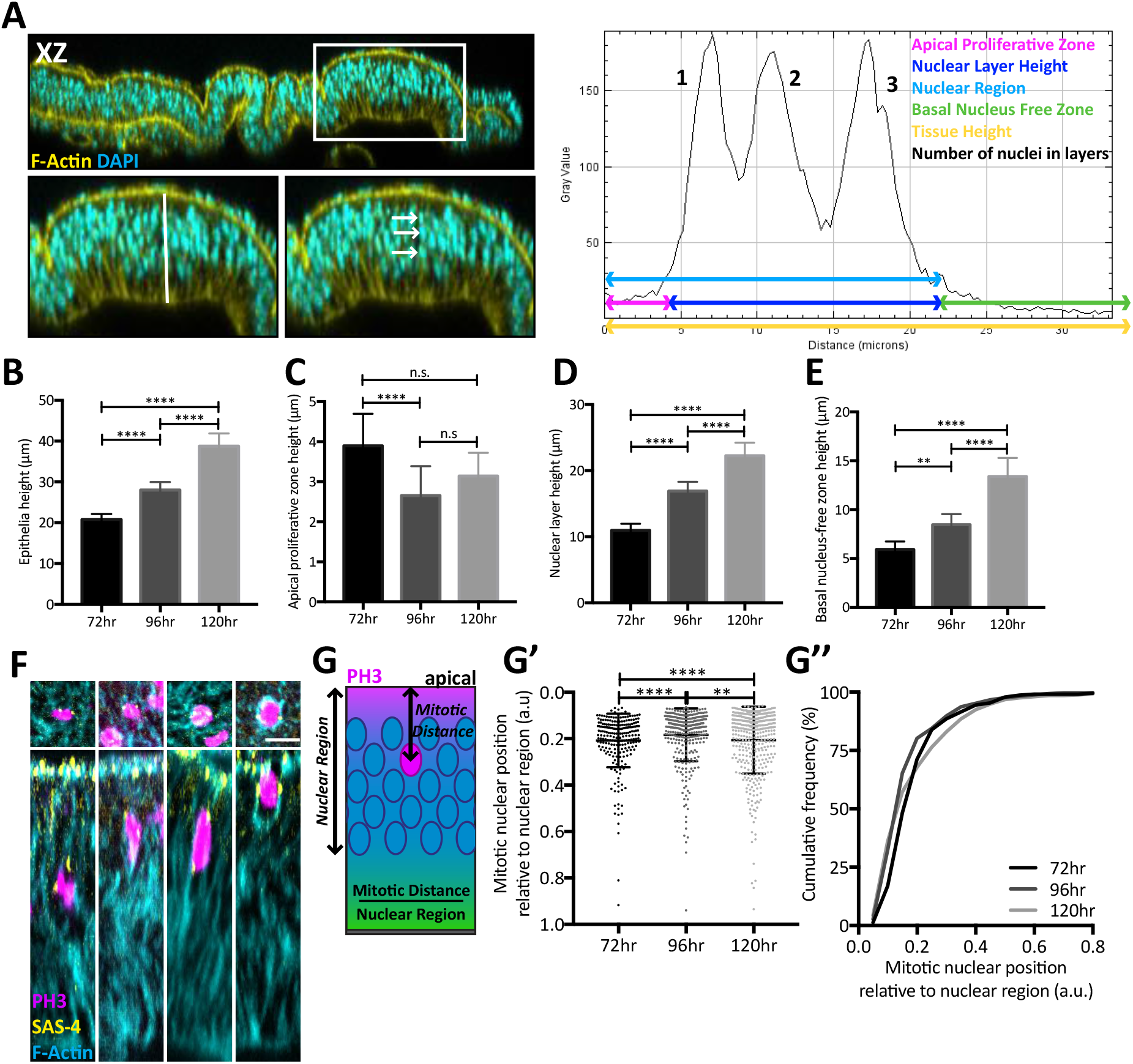
Changes in the wing disc architecture during development are associated with distinct patterns of mitotic nuclear positioning. (A) Left panel, lateral cross section of wing disc stained with DAPI (cyan) and phalloidin (yellow). White box highlights inset pouch region shown below. White line in lower left inset highlights the apico-basal cross-section used to generate the intensity plot displayed in the right panel. Lower right inset: white arrows mark the nuclei shown as peaks in the right panel, used for counting numbers of nuclei in the nuclear layers. Right panel: labelled intensity plot illustrating how measurements were obtained for Figure 1B’,C and S1B-E. (B-E) Measurements of apico-basal architectural features at distinct developmental stages of the wing disc, the same data are presented in a condensed manner in Figure 1B’. (F) Sections of wing disc pouch region stained with anti-PH3 (magenta), anti-SAS-4 (yellow) and phalloidin (Cyan) to mark mitotic cells, centrosomes and actin, respectively. Top view shows single planar cross section. Lower view shows lateral cross section. Outside of mitosis, Sas-4 marked centrosomes localise to the apical surface, while mitotic centrosomes localise to the nucleus at varying distances apart. We observed centrosome arrangements consistent with mitosis, independent of the distance of PH3+ nuclei from the apical surface confirming these nuclei are mitotic. As mitosis only occurs at the apical surface in wildtype wing discs, this indicates these are mitotic nuclei captured on their apical trajectory. (G) Schematic showing measurement of the mitotic nuclear position relative to the nuclear region. (G’) PH3+ nuclear distance shown in Figure 1E-E’’ normalised to the height of the nuclear region to give the relative position within the region occupied by nuclei. Average height of nuclear region calculated per disc. (G’’) Data in G’ presented as a cumulative frequency distribution. (B-E and G) n = 8, 8 and 5 wing discs for 72, 96 and 120hr respectively. Scale bar in F is 5*µ*m. Statistical significance in B-E are given for one-way ANOVA. Statistical significance in G given for Kolmogorov-Smirnov comparison of cumulative distribution. n.s. > 0.05, **p < 0.01, ****p < 0.0001. Error bars represent mean ± SD in all plots.

**Figure S2, Supplementary data for Figure 2:**
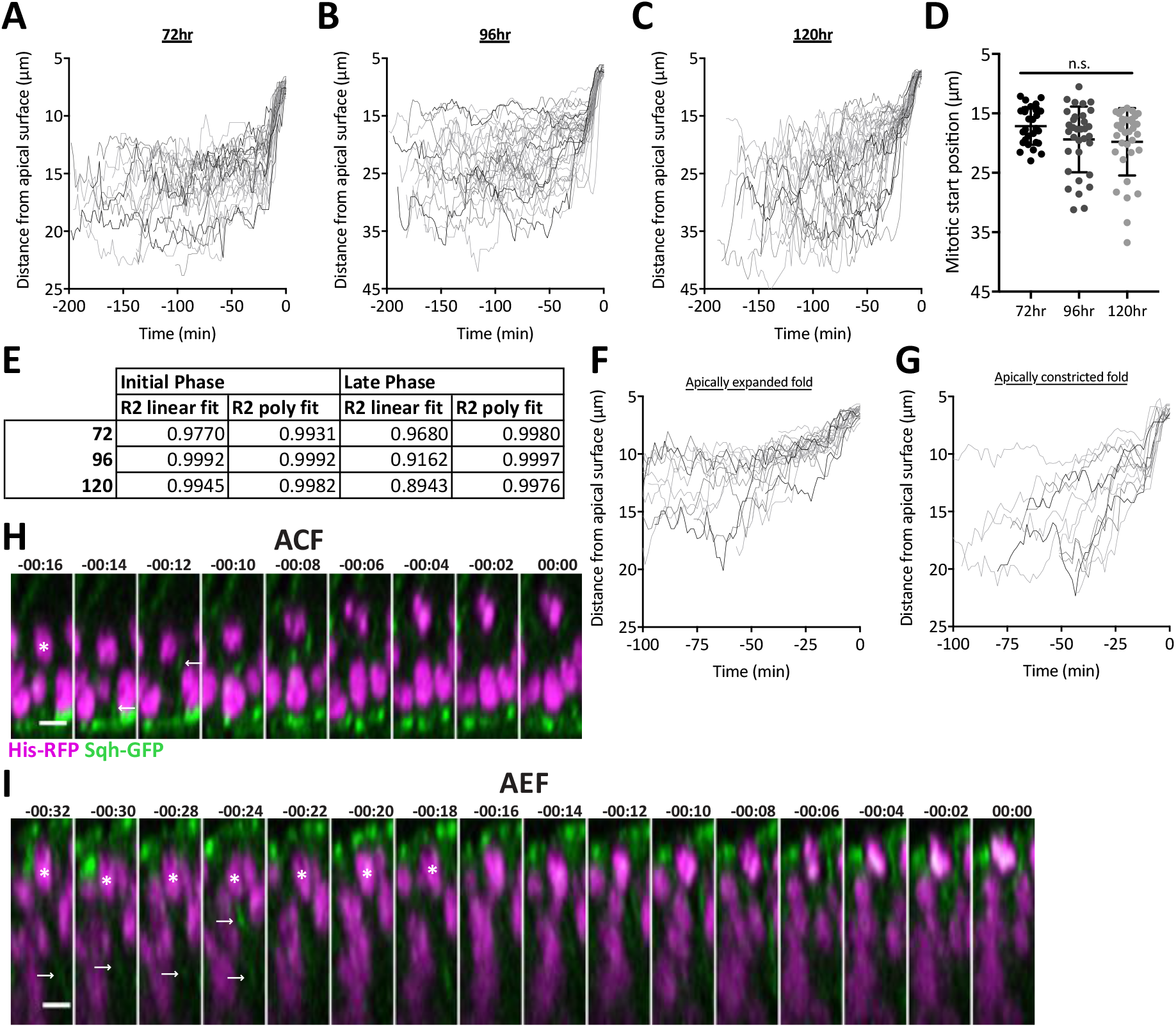
Apical nuclear movement is driven by mitotic rounding and nuclear dynamics depend on tissue architecture. (A-C) Individual nuclei tracks from the wing disc pouch region at 72, 96 and 120 hours AEL respectively, expressing Sqh-GFP and His RFP, obtained by measuring distances illustrated in Figure 2F. 0 minutes is defined by metaphase nucleus prior to anaphase separation. (D) The position at the onset of apical nuclear movement, used to calculate mitotic nuclear velocity in Figure 2E. (E) Mean square displacement R^2^ values for linear and polynomial fits for early and late phase of nuclear tracking, as shown in Figure 2B’-D’ at wing disc developmental stages. (F-G) Individual nuclei tracks from the wing disc fold regions described in Figure 2G and averaged in Figure 2J. (H-I) Representative time-lapses of mitotic nuclei moving apically in the distinct fold regions in wing discs expressing Sqh-GFP and His-RFP. White arrows indicate cortical myosin enrichment. White asterisks mark the tracked nucleus when in crowded regions. (A-E) n = 28, 33 and 34 nuclear trajectories from 3, 3 and 4 wing discs for 72, 96 and 120 hr respectively. (F) n = 14 from 3 wing discs. (G) n = 20 nuclear trajectories from 3 wing discs. Scale bars in H-I are 5 *µ*m. Statistical significance in D is given for one-way ANOVA. n.s. p > 0.05. Error bars represent mean ± SD in D.

**Figure S3, Supplementary data for Figure 3:**
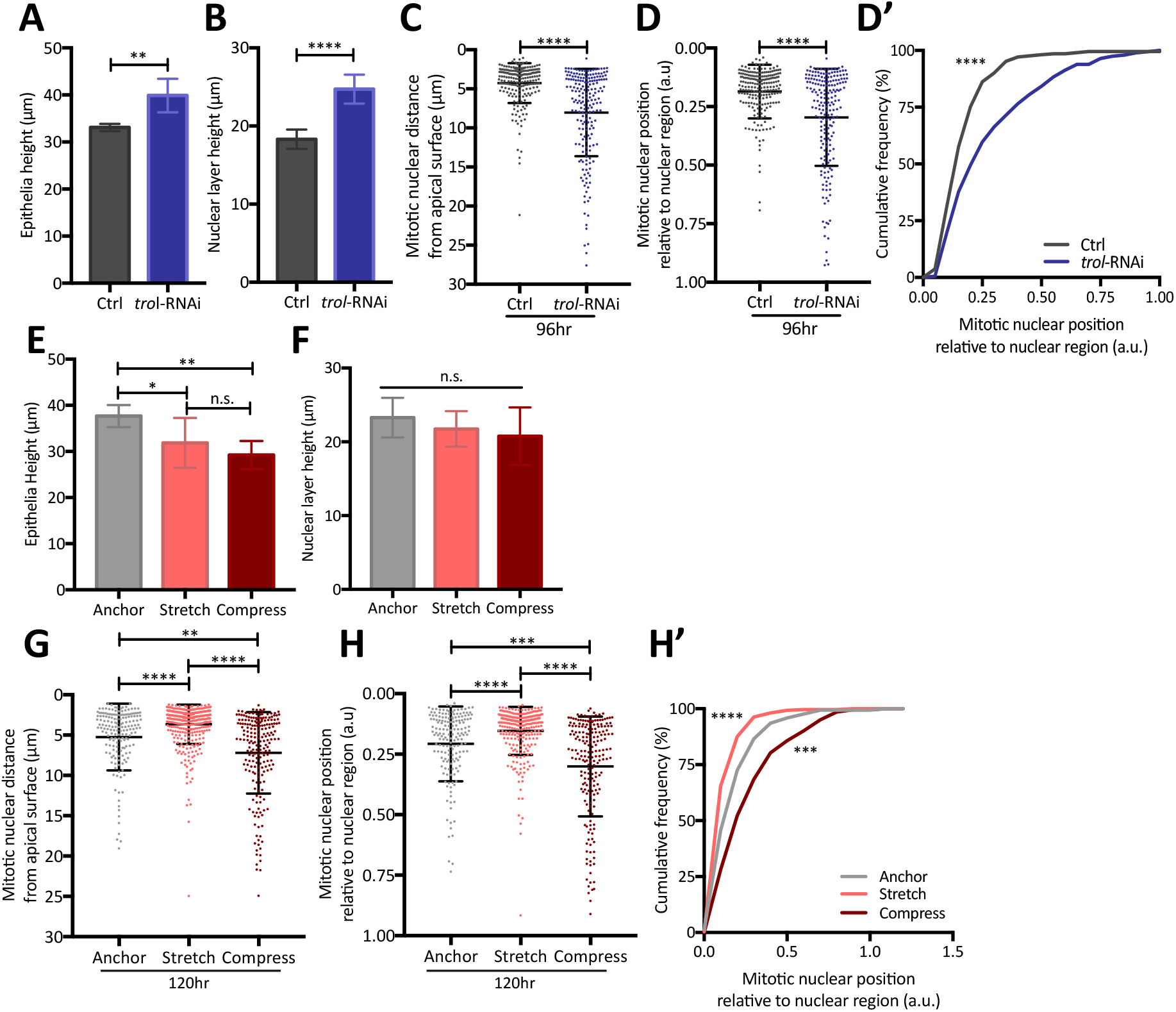
Perturbing cell density influences mitotic nuclear positioning. (A-B) Quantification of apico-basal epithelial morphology for control (*actin*Gal4) and U*AS-trol*-RNAi wing discs. Data corresponding to Figure 3B. (C) Scatter plot of mitotic nuclear distance from the apical surface for control (*actin*Gal4) and U*AS-trol*-RNAi wing discs. Data corresponding to Figure 3D. (D) Scatter plot of mitotic nuclear position, relative to the nuclear region for control (*actin*Gal4) and U*AS-trol*-RNAi expressing wing discs. (D’) Cumulative frequency distribution for data in D. (D-D’) Data corresponding to Figure 3D and S3C. (E-F) Quantification of apico-basal epithelial morphology in the wing disc pouch region upon mechanical perturbation as illustrated in Figure 3E. (G) Scatter plot of mitotic nuclear distance from the apical surface upon mechanical perturbation. Data corresponding to Figure 3I. (H) Scatter plot of mitotic nuclear position, relative to the nuclear region in wing disc pouch region, upon mechanical perturbation. (H’) Data in H presented as cumulative frequency distribution. (H-H’) Data corresponds to Figure 3I and S3G. (A-D’) n = 6 and 7 wing discs for control and *trol*-RNAi respectively. (E-H’) n = 3, 8 and 6 wing discs for anchor, stretch and compress conditions respectively. Statistical significance given for unpaired t-test in A-B, Kolmogorov-Smirnov comparison of cumulative distribution in C-D’ and G-H’, and one-way ANOVA in E-F. *p < 0.05, **p < 0.01, ***p < 0.001, ****p < 0.0001. Error bars represent mean ± SD in all plots.

**Figure S4, Supplementary data for Figure 4:**
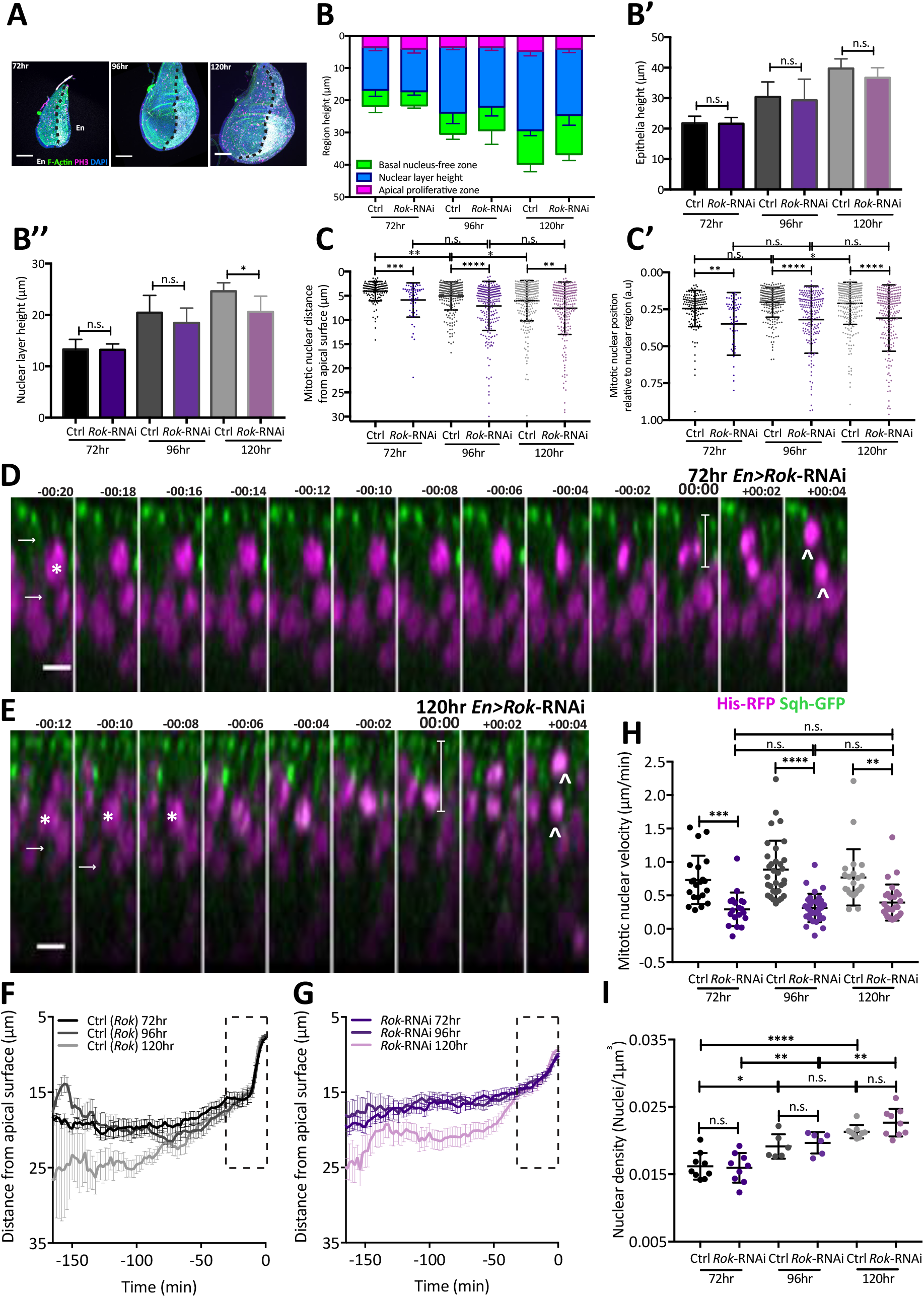
Rok is required for apical nuclear movement at all stages of development. (A) Representative images of wing disc morphology with *Rok*-RNAi in the *engrailed* domain (posterior, *en*) at different developmental stages, marked by UAS-NLS-Cherry in the posterior region and phalloidin (green), anti-PH3 (magenta) and DAPI (blue). Black dashed line shows anterior-posterior boundary. Image accompanies Figure 4A. (B-B’’) Measurements of epithelial apico-basal morphology in anterior control region and posterior region expressing *Rok*-RNAi. (C) Scatter plot of absolute mitotic nuclear distance from apical surface for control (anterior) and *Rok*-RNAi (posterior) wing disc regions at different developmental stages, corresponds to Figure 4B. (C’) Scatter plot of mitotic nuclear position, relative to the nuclear region for anterior control and posterior *Rok*-RNAi expressing wing disc regions at different developmental stages, corresponds to Figure 4B’. (D-E) Representative time-lapse of mitotic nuclei moving apically within *Rok*-RNAi expressing, posterior wing pouch region in a background of Sqh-GFP and His-RFP, at 72 (D) and 120 (E) hours AEL. White arrows indicate cortical myosin enrichment regions. White asterisks mark the tracked nuclei when in crowded regions. The last two time points show failure to orient planar cell division, daughter chromosomes marked by arrow-heads. White line shows distance of metaphase nucleus from apical surface at 0 minutes, with values presented in Figure 4E. (F-G) Full time course for average nuclear tracks for anterior control (F) and posterior *Rok-*RNAi (G) mitotic cells, at each developmental stage. Black dashed line represents region enlarged in Figure 4D-D’’’. (H) Mitotic nuclear velocity for control and *Rok*-RNAi expressing mitotic cells, at each developmental stage. Each dot represents a single tracked nucleus. Calculated from onset of apical movement and cortical mitotic enrichment, up to most-apical position, i.e. metaphase. (I) Nuclear density per *µ*m^3^ for anterior control and posterior *Rok*-RNAi expressing wing disc regions, in SqhGFP and HisRFP backgrounds, at different developmental stages. (B-C’) n = 8, 7 and 5 wing discs for 72, 96 and 120 hours respectively (F-H) n = 19-33 nuclear trajectories acquired from 3 wing discs per developmental stage. Scale bars: A, 50 *µ*m, D-E, 5 *µ*m. Statistical significance given for unpaired t-test in B’-B’’, Kolmogorov-Smirnov comparison of cumulative distribution in C-C’ and one-way ANOVA in H. n.s. p > 0.05, *p < 0.05, **p < 0.01, ***p < 0.001, ****p < 0.0001. Error bars represent mean ± SEM in F-G and mean ± SD in B-C’, H and I.

**Figure S5, Supplementary data for Figure 5:**
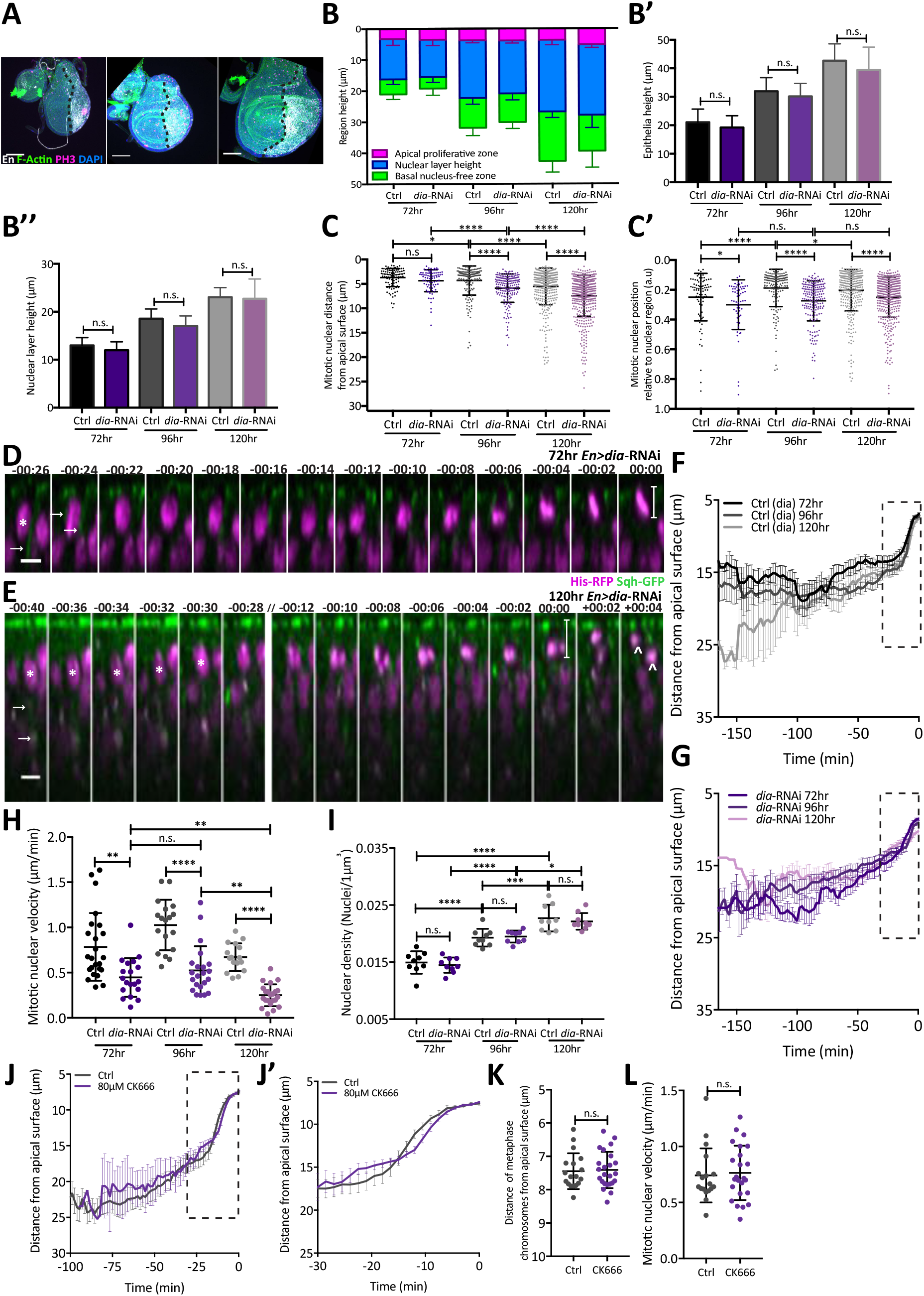
Dependency on Dia for apical mitotic nuclear movement increases with development. (A) Representative images of wing disc morphology with *dia*-RNAi in the *engrailed* domain (posterior, *en*) at different developmental stages, marked by UAS-NLS-Cherry in the posterior region and phalloidin (green), anti-PH3 (magenta) and DAPI (blue). Black dashed line shows anterior-posterior boundary. Image accompanies Figure A. (B-B’’) Measurements of epithelial apico-basal morphology in anterior control region and posterior region expressing *dia*-RNAi. (C) Scatter plot of absolute mitotic nuclear distance from apical surface for control (anterior) and *dia*-RNAi (posterior) wing disc regions at different developmental stages corresponds to Figure 5B. (C’) Scatter plot of mitotic nuclear position, relative to the nuclear region for anterior control and posterior *dia*-RNAi expressing wing disc regions at different developmental stages, corresponds to Figure 5B’. (D-E) Representative time-lapse of mitotic nuclei moving apically within *dia*-RNAi expressing, posterior wing pouch region in a background of Sqh-GFP and His-RFP, at 72 (D) and 120 (E) hours AEL. White arrows indicate cortical myosin enrichment regions. White asterisks mark the tracked nuclei when in crowded regions. The last two time points show failure to orient planar cell division, daughter chromosomes marked by arrow-heads. White line shows distance of metaphase chromosomes from apical surface at 0 minutes, with values presented in Figure 5E. (F-G) Full time course for average nuclear tracks for anterior control (F) and posterior *dia-*RNAi (G) mitotic cells, at each developmental stage. Black dashed line represents region enlarged in Figure 5D-D’’’. (H) Mitotic nuclear velocity for control and *dia*-RNAi expressing mitotic cells, at each developmental stage. Each dot represents a single tracked nucleus. Calculated from onset of apical movement and cortical mitotic enrichment, up to most-apical position, i.e. metaphase. (I) Nuclear density per *µ*m^3^ for anterior control and posterior *dia*-RNAi expressing wing disc regions, in SqhGFP and HisRFP backgrounds, at different developmental stages. (J-J’) Average nuclear tracks for control and CK666 treated wing discs. Black dashed line represents region enlarged in I’. (K) The final distance of the metaphase chromosomes from the apical surface for control and CK666 treated wing discs. (L) Mitotic nuclear velocity for control and CK666 treated wing discs. Each dot represents single tracked nucleus. Calculated from onset of apical movement up to most-apical position, i.e. metaphase. (B-C’) n = 4, 7 and 8 wing discs for 72, 96 and 120 hours respectively (F-H) n = 16-24 nuclear trajectories acquired from 3-4 wing discs per developmental stage. (I-K) n = 18 and 24 nuclear trajectories for acquired from 3 control and 3 CK666 treated wing discs respectively. Scale bars: A, 50*µ*m, D-E, 5*µ*m. Statistical significance given for unpaired t-test in B’-B’’, J and K, Kolmogorov-Smirnov comparison of cumulative distribution in C-C’ and one-way ANOVA in H. n.s. p > 0.05, *p < 0.05, **p < 0.01, ****p < 0.0001. Error bars represent mean ± SEM in F-G, J-J’ and mean ± SD in B-C’, H, I, K and L.

**Figure S6, Supplementary data for Figure 6:**
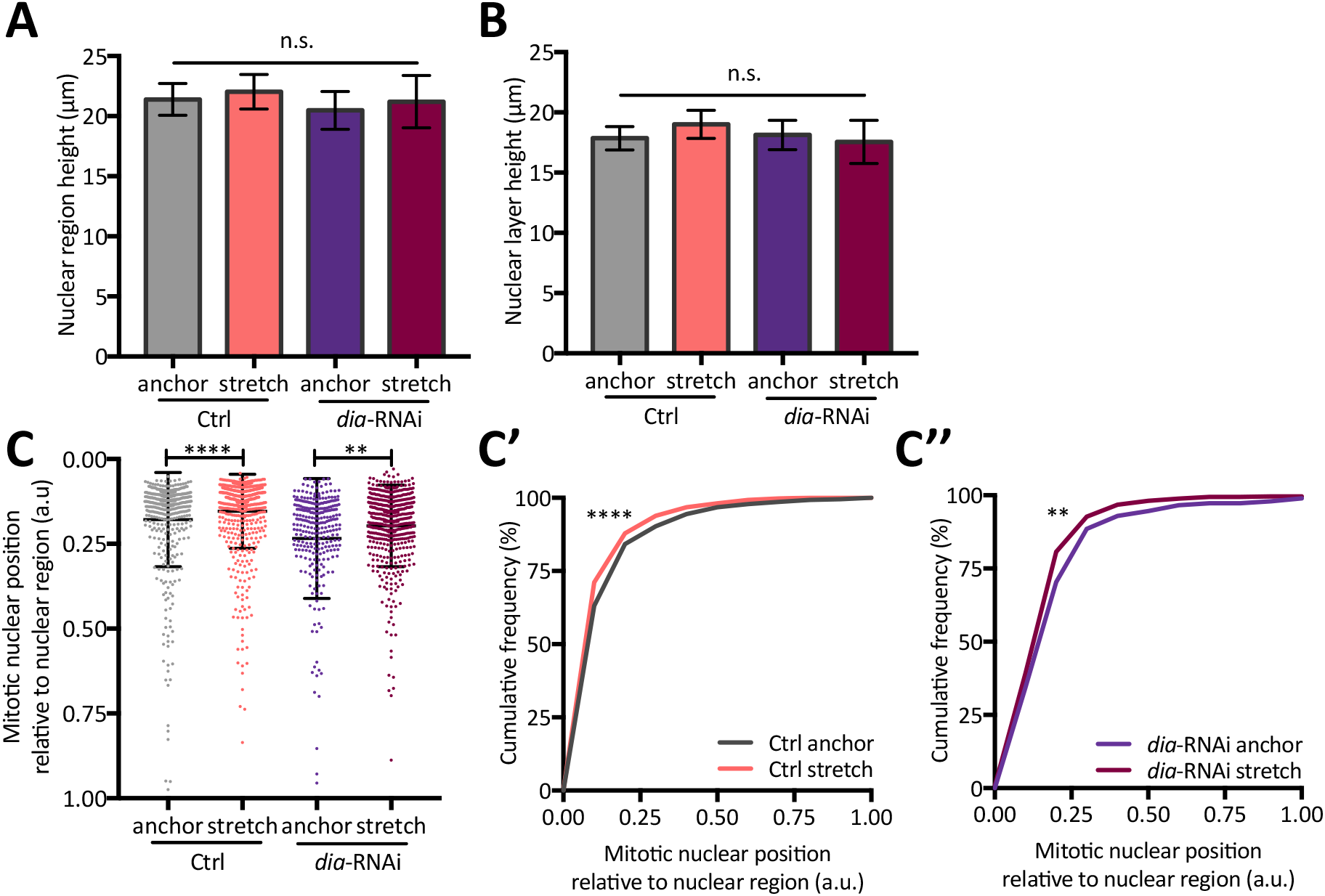
Defects in nuclear positioning upon Dia depletion can be rescued by mechanically reducing cell density. (A-B) Measurements of epithelial apico-basal morphology for control and *dia*-RNAi wing disc pouch region, in anchor and stretched conditions (C) Mitotic nuclear position relative to the nuclear region for control and *dia*-RNAi expressing wing disc pouch regions in anchor and stretched conditions. (C’-C’’) Data shown in C presented as cumulative frequency distribution, control (C’) and *dia*-RNAi (C’’). Data corresponds to Figure 6C-C’’ and S6C. (A-C’’) n = 6 and 6 wing discs for anchor and stretch, control conditions respectively and n = 7 wing discs each for *dia*-RNAi anchor and stretch. Statistical significance given for one-way ANOVA in A-B, Kolmogorov-Smirnov comparison of cumulative distribution in C-C’’. n.s. p > 0.05, **p < 0.01, ****p < 0.0001. Error bars represent mean ± SD in A-C.

